# Site-specific phosphorylation of Ser352 drives aggregation of Tau R4 under acidosis conditions

**DOI:** 10.64898/2026.05.03.722553

**Authors:** Shachar Guy Bressler, Carmia Blacher, Karina Abramov Harpaz, Dana Grunhaus, Stefan G.D. Rüdiger, Mattan Hurevich, Deborah E. Shalev, Yifat Miller, Assaf Friedler

**Affiliations:** The Institute of Chemistry, The Hebrew University of Jerusalem, Edmond J. Safra Campus, Givat Ram, Jerusalem, 91904, Israel; Department of Chemistry, Ben-Gurion University of the Negev, P.O. Box 653, Be’er Sheva 8410501, Israel; Ilse Katz Institute for Nanoscale Science and Technology, Ben-Gurion University of the Negev, Beér-Sheva 8410501, Israel; The School of Brain Sciences and Cognition, Ben-Gurion University of the Negev, Beér-Sheva 8410501, Israel; Protein Chemistry of Disease, Department of Chemistry, Utrecht University, Padualaan 8, 3584CH, Utrecht, The Netherlands; Cellular Protein Chemistry, Bijvoet Center for Biomolecular Research, Utrecht University, Padualaan, 8, 3584CH Utrecht, The Netherlands; Science for Life, Utrecht University, Padualaan 8, 3584CH Utrecht, The Netherlands; Department of Pharmaceutical Engineering, Azrieli College of Engineering Jerusalem, Jerusalem 91035, Israel

## Abstract

In Alzheimer’s disease, neurons undergo acidosis as their pH drops from ∼7.1 to ∼6.5. Here we elucidate the molecular mechanism of specific Tau aggregation only at this lower pH. We show that a specific phosphorylation event in the Tau R4 domain is coupled to this pH drop to drive a defined phase transition from condensates to fibrils. Using a combination of experimental and computational studies, our results demonstrate that site-specific phosphorylation of Ser352 is the molecular switch that induces aggregation of Tau only at the more acidic, disease-related pH. We designed and synthesized a phosphopeptide library covering all phosphorylation patterns of the Tau-R4 domain and tested its response to controlled acidification using an array of complementary biophysical methods. This revealed that only a single phosphorylation of Ser352 led to acid-driven aggregation of Tau-R4. At neutral pH, Tau-R4 with a phosphorylation on Ser352 formed liquid condensates. These condensates converted irreversibly into amyloid filaments as the pH gradually decreased towards the level associated with pathological acidosis. Using a combination of ³¹P-and ^1^H - NMR, MD simulations and microscopy studies, we show that the mechanism by which the phosphorylation of Ser352 Tau R4 exerts its effects is based on the unique position of this residue within the protein structure. pSer352 is located at the inside of the tip of a β-hairpin, pointing into the hydrophobic core of the amyloid fold. This buried position increases its effective pK□, resulting in compacting of the hairpin following pSer352 protonation upon acidification. This shortens the inter-sheet distance at this position and tightens the filament. We conclude that this single protonation event of the pSer352 phosphate is responsible for the macroscopic phase transition from condensates to aggregates. Our results provide the molecular explanation for the specific aggregation of Tau under the pathologically relevant acidic pH, which is substantially different from its behavior at neutral pH.

## INTRODUCTION

Tau is a microtubule-binding protein that stabilizes microtubules (MT) in neurons. Hyperphosphorylation of Tau followed by its aggregation is implicated in neurodegenerative diseases (ND) such as Alzheimer’s (AD)^1^. Tau binds MT via its microtubule binding region, which contains four repeats (R1-R4), and a pseudo-R domain termed R’ (Figure 1a). Hyper-phosphorylated Tau dissociates from microtubules and forms amyloid fibrils, a hallmark of AD^2^. ND are also associated with lower intracellular pH values, caused by impaired lysosomal acidification due to decreased v-ATPase activity^3^, in a phenomenon termed acidosis^4^. Acidification may also promote Tau pathology indirectly by activating asparaginyl endopeptidase, a lysosomal protease whose activity is favored under acidic conditions. AEP activation in Alzheimer’s disease is linked to increased Tau hyperphosphorylation, through cleavage of the PP2A inhibitor SET, resulting in suppression of PP2A activity and shifting Tau toward a hyperphosphorylated state^5,6^. In Alzheimer’s disease patients, the pH in neurons decreases from the physiological range of 7.3-7.1 to approximately 6.5^7–12^. Acidosis drives the aggregation of proteins that are associated with ND. For example, the half-time of α-synuclein aggregation at pH 7 is approximately 80 h, while at pH 6 the aggregation accelerates by almost 20-fold, to an average of 5 h^13^. Amyloid beta (Aβ) shows a similar trend: At pH 7.1 its aggregation half-time is 15 h, while at pH 6.5 aggregation occurs in approximately 2 h^14^. pH decrease is also a key factor in the aggregation of Tau: Non-phosphorylated Tau 297-391 was found to form filaments at pH values of 6 and 5^15^.

**Figure 1.**
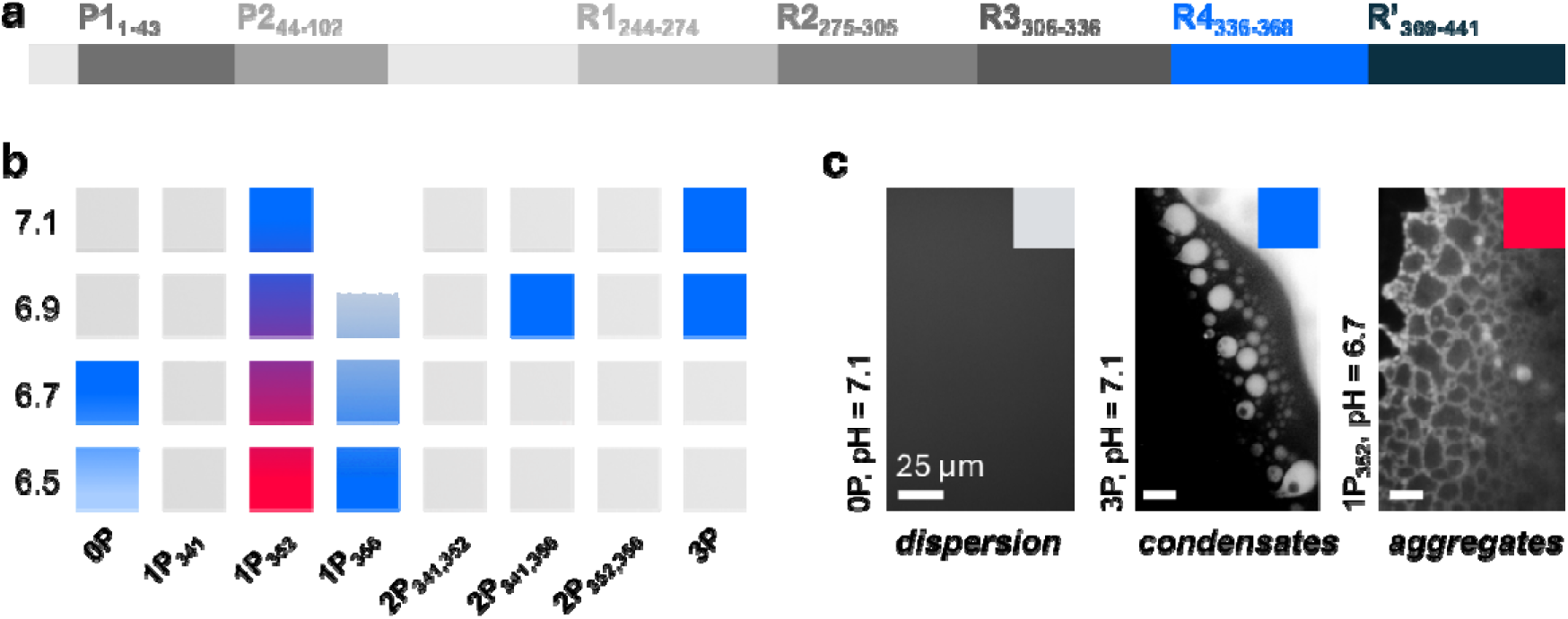
Condensation and Aggregation of 1P_352_ is pH dependent. a) Domain structure of Tau. b) Fluorocsence microscopy experiments show the condensation and aggregation properties of the Tau-R4 library at different pH values. Blue squares indicate condensation, red squares indicate aggregation and a mixture of the two colors represents that both processes took place. Grey indicates no observed condensation or aggregation. c) Representative fluoroscence microscopy images showing 0P as an example of soluble peptides (left), 3P as an example of condensates (middle), and 1P_352_ as an example of aggregates (right).

Condensation of Tau, a process that leads to formation of droplets via liquid-liquid phase separation (LLPS), is a crucial step in its aggregation mechanism^16^, although the molecular mechanism is not yet fully understood. Studies at neutral pH show that aggregation and condensation of microtubule associated proteins (MAP), particularly Tau, are strongly dependent on specific phosphorylation patterns^17–23^. For example, incubation of Tau with a mixture of the SPAK4 and PKA kinases led to aggregation of Tau. However, incubation of Tau with a mixture of GSK-3β and PKA, that generate different phosphorylation patterns did not^19^. Single phosphorylation of serines 293 or 305 led to aggregation of Tau^24^. We have previously shown that the phosphorylation pattern determines whether the R4 domain of Tau (Tau 336-358, Tau-R4) undergoes condensation, aggregation, or remains soluble^25^. Tau-R4 forms a conserved hairpin structure in many ND-derived amyloid filaments, including chronic traumatic encephalopathy and AD^26^. Specifically, we found that the phosphorylation of Ser341 promotes aggregation at pH 7.4, while phosphorylation of Ser352 leads to condensation, and Ser356 phosphorylation did not induce condensation or aggregation^25^.

To determine whether the mildly acidic pH environment found in brains of neurodegenerative diseases patients selectively affects Tau aggregation in a phosphorylation dependent manner, we systematically examined how a pH decrease from 7.1 to 6.5 affects the condensation and aggregation of Tau-R4 across all possible phosphorylation patterns of its three serine residues. Our results identified the single phosphorylation of Ser352 as the driving force for aggregation at the clinically-relevant acidic pH.

## RESULTS

### Screening a library of the R4 phosphorylation patterns revealed that 1P_352_ undergoes condensation-mediated aggregation upon decrease in pH

Our previous results have shown that serine phosphorylation plays a key role in regulating the aggregation of Tau-R4 at physiological pH ^25^. We use synthetic peptides corresponding to the full R4 domain as a model system that enables the construction of a library comprising all the specific phosphorylation patterns with full control over stoichiometry and position^27^. Phosphoserine has a theoretical pKa value in the range of 5.6^28^-6.1^29,30^, which means it can change its protonation state around the pathological pH value of 6.5. Here we tested how a pH decreases shifts Tau-R4 from condensation to aggregation, and how different phosphorylation patterns regulate this response. To achieve this, we synthesized a Tau-R4 (336-358) peptide library comprising all possible phosphorylation patterns^27^ (Table 1) and tested its aggregation and condensation properties using fluorescence microscopy at a pH range starting at the physiological pH value of 7.1 and gradually decreasing to pH 6.5. The pH value of 6.5 was selected as the end point of the titration since this is the pH value observed in the brains of Alzheimer’s disease patients^7–12^.

**Table 1.**
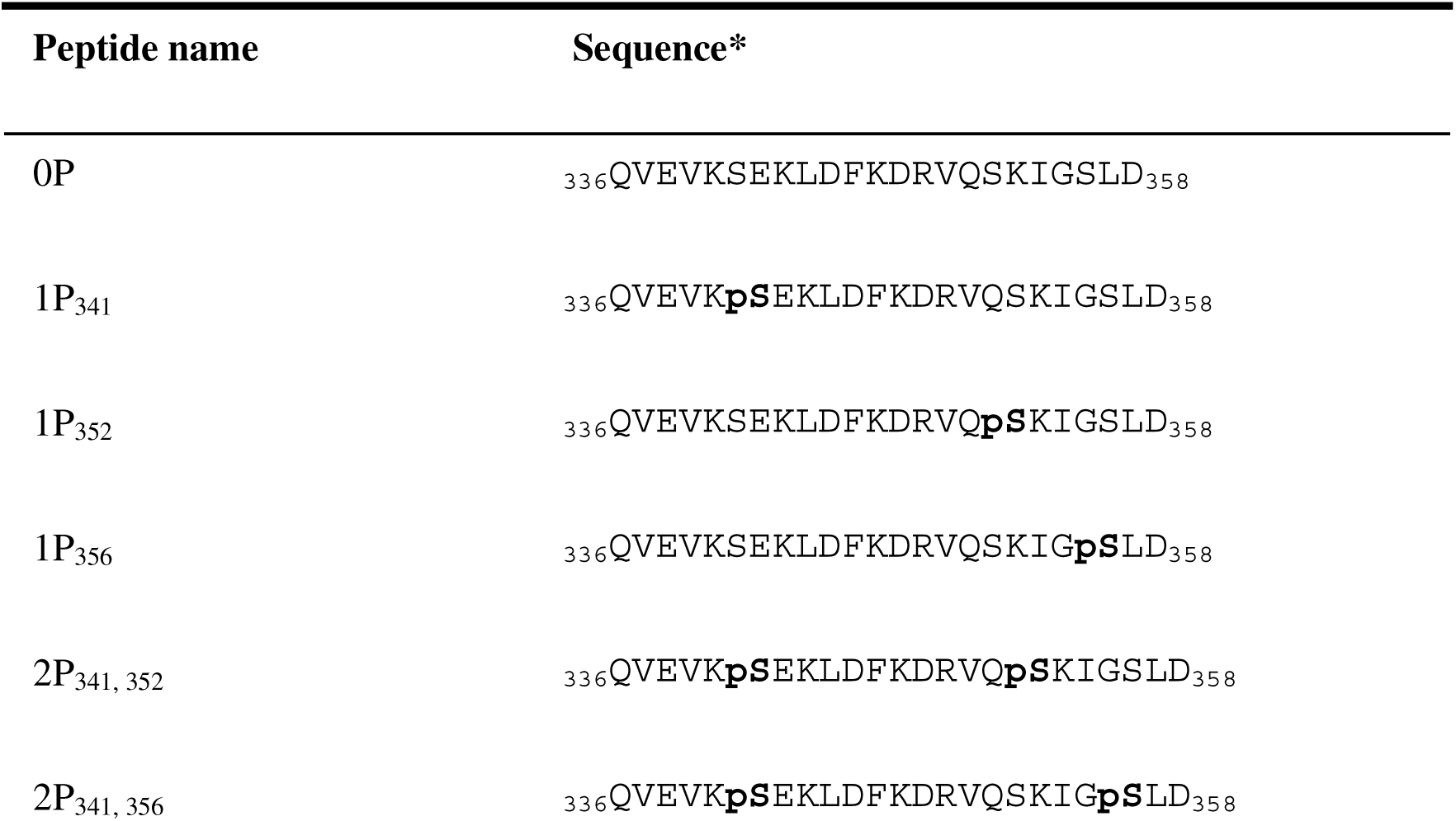
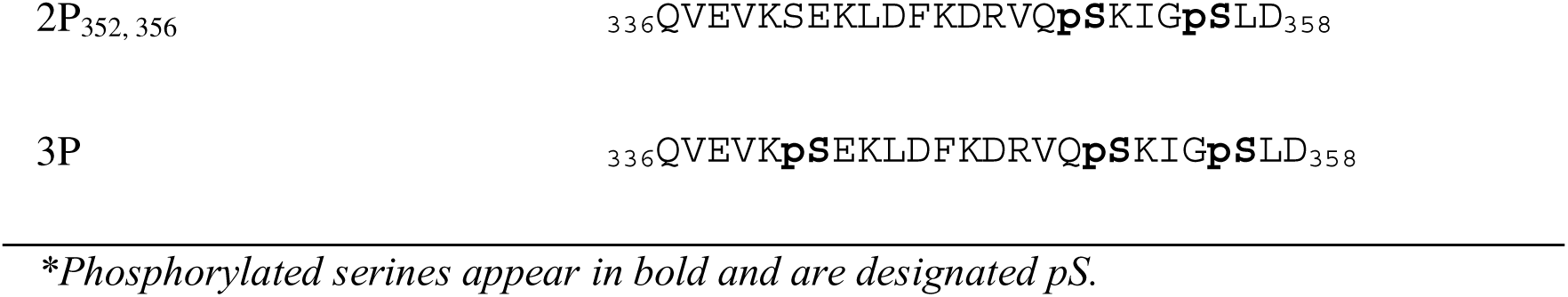
Tau-R4 (336-358) and its derived phosphorylated peptides used in this study.

The non-phosphorylated R4 domain between residues Q336-D358 (termed here Tau-R4 or 0P), formed condensates only under the more acidic conditions of pH 6.7 and 6.5 (Figure 1b, c) and did not aggregate at pH values of 7.1 and 6.5. The mono-phosphorylated R4 peptide 1P_352_ underwent condensation at pH 7.1 and 6.9. It formed aggregates at lower pH values of 6.7 and 6.5. At pH 6.7, we found coexistence of droplets and aggregates with aggregation predominating (Figure 1b, c), while at 6.5 we found only string-like aggregates. This indicates that pH 6.9 to 6.7 is the transition range between condensation and aggregation of 1P_352_. 1P_356_ formed droplets at pH values of 7.1 and 6.9. 1P_341_ did not undergo self-assembly at any pH. Among the multi-phosphorylated R4 peptides, we found that 2P_341,352_, and 2P_352,356_ showed neither condensation nor aggregation across all tested pH values. 2P_341,356_ and the fully phosphorylated domain 3P underwent condensation at pH 6.9, with Tau-R4_(3P)_ also forming condensates at the neutral pH of 7.1. These findings highlight pSer352 as a specific phosphorylation that can initiate aggregation under acidic conditions, emphasizing its potential role in the pathogenesis of AD.

### 1P352 undergoes condensation-mediated aggregation upon decrease in pH

To study the mechanism of aggregation at pH = 6.5, we titrated 1P_352_ in a buffer solution from pH 7.1 to 6.5, decreasing the pH by 0.1 units each time, and monitored the droplets using fluorescence microscopy. At pH values of 7.1, 7.0 and 6.9 only droplets were obtained. Decreasing the pH to 6.8 resulted in droplets that were smaller in diameter, with some of them agglomerating to form aggregates (Figure 2a). At pH 6.7 we observed the formation of glass-like conglomerated droplets. When the pH was further decreased to 6.6 and 6.5, tighter arrangements of droplets were observed. Using the ImageJ 1.54j software, we analyzed the cross-section of the droplet profile. The analysis showed that lower pH values led to smaller droplets that formed a thinner assembly overall. (Figure 2b, 2c).

**Figure 2.**
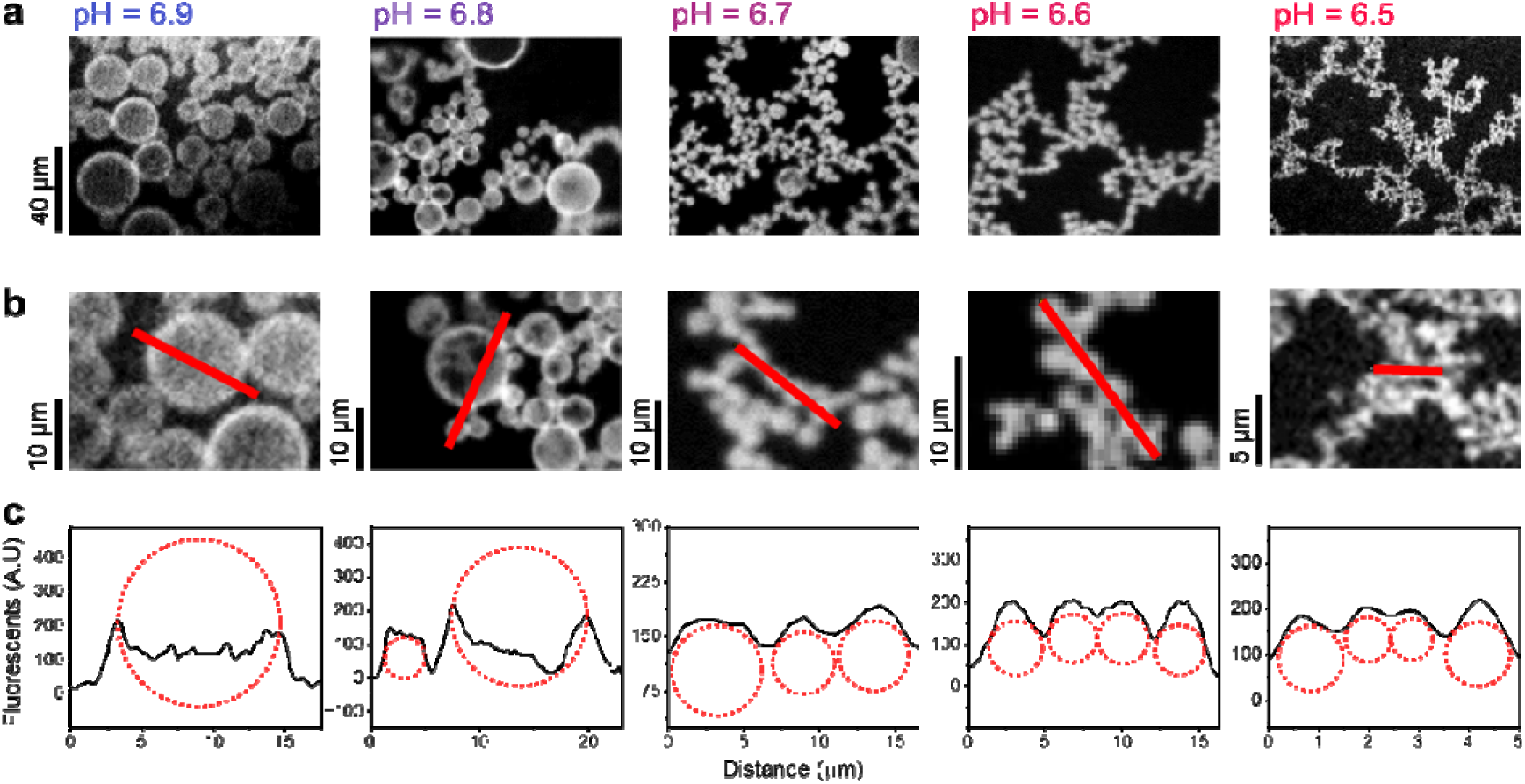
1P_352_ aggregates via condensation at lower pH values. a) Fluorescent microscopy images of 1P_352_ at pH 6.9, 6.8, 6.7, 6.6 and 6.5. b) Expanded view of the droplets, with a red line highlighting the corresponding cross-section shown in panel c. c) Cross-section analysis of the droplets at all pH values tested. The locations of the droplets are shown as red circles.

Analysis of the effect of the pH on the size of the droplets showed that the diameter of the droplets decreased from an average of 7.1 ± 0.5 µm at pH 7.0, to 5.8 ± 0.4 µm at pH 6.9, and further decreased to 4.1 ± 0.7 µm at pH 6.8. Further pH decreases from 6.8 to 6.7 led to an average diameter of 2.4 ± 0.1 µm. When going from pH 6.7 to 6.5, the overall change was to a final diameter of 0.73 ± 0.04 µm (Figure 3a). The critical pH value for this transition, calculated by a sigmoidal fit to the diameter of the droplets, was found to be 6.80 ± 0.02. These results suggest that upon phosphorylation of Ser352 in Tau, small changes of ∼0.3 pH units from normal neurons can be crucial for the initiation of an aggregation process.

**Figure 3.**
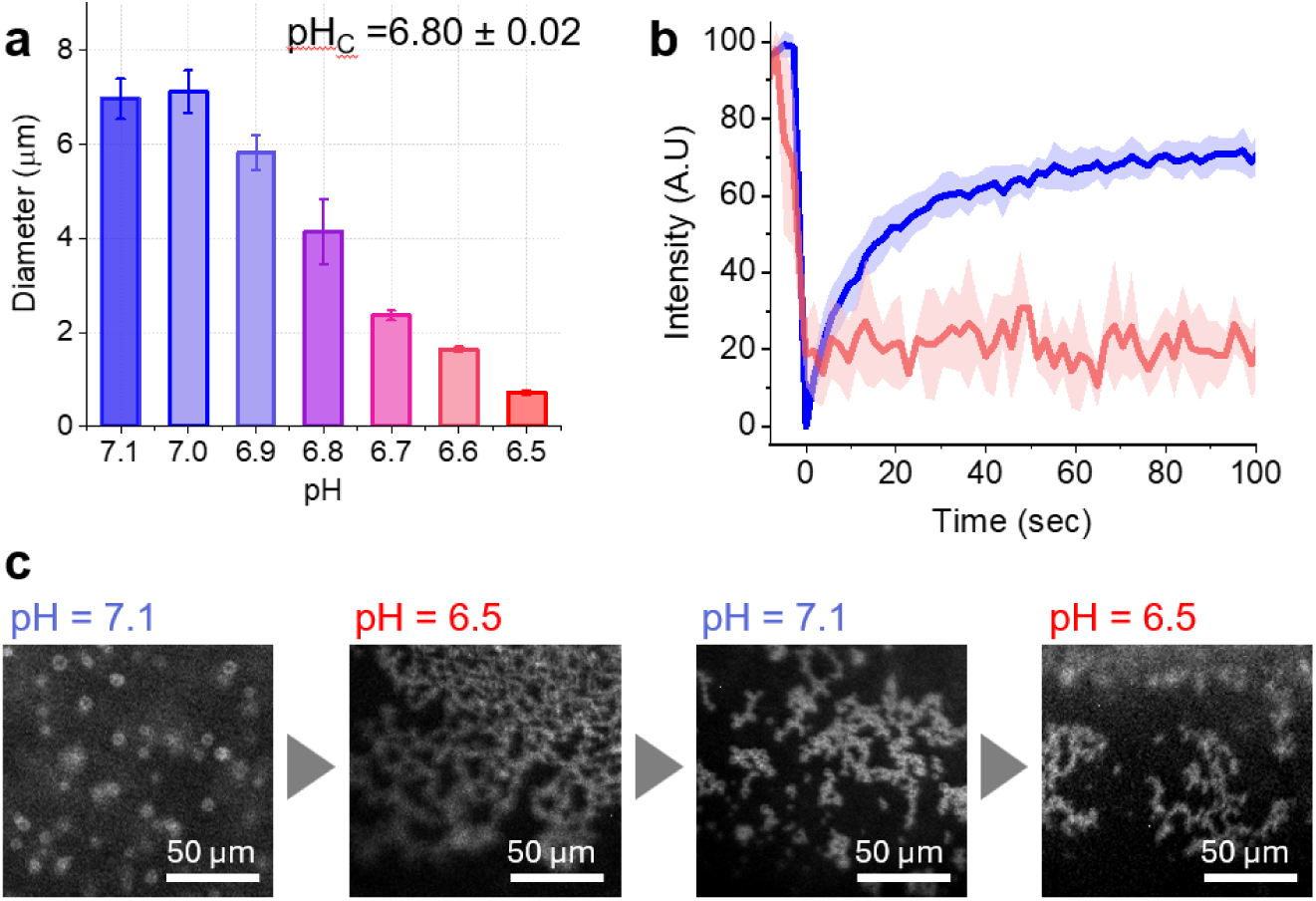
Upon decreasing the pH, the size of the 1P_352_ droplets decreased and the droplets formed thermodynamically stable aggregates. a) Average diameters of the droplets formed by 1P_352_ show that the lower the pH, the smaller the droplets are. b) Fluorescence recovery after photobleaching (FRAP) plot of 1P_352_ at pH 7.1 (blue) and 6.5 (red). c) Stability assay shows that after the first pH decrease, the aggregates are the predominant species, suggesting that aggregation is an irreversible process.

Condensates exchange peptides with the surroundings due to diffusion, and thus exhbit fast recovery after photobleaching. Aggregates, on the other hand, do not^16^. To characterize the effect of pH on the diffusion rate in the droplets, we determined the diffusion coefficients of the droplets using fluoroescent recovery after photobleaching (FRAP). We found that the half time for recovery at pH 7.1 was 2 ± 1 s and the diffussion coefficient was 3.3×10^-12^m^2^/s. At pH 7.1, the droplets of 1P_352_ reached a recovery of 75 % of the initial intensity (Figure 3b), suggesting 25 % of immobile peptide within the condensates. In contrast, at pH of 6.5 the aggregates showed no recovery at all. Hence, we concluded that decreasing the pH turns the droplets into more solid spheres that agglomerate to form glass-like condensates.

### 1P352 filaments do not dissociate to condensates upon increasing the pH from 6.5 to 7.1

The stability of the aggregates once formed was assessed by gradually increasing the pH back to 7.1 and monitoring by fluorescence microscopy whether the aggregates dissolved and condensates reformed. Aggregates formed at pH 6.5 remained unchanged and did not revert to condensates upon increasing the pH to 7.1 (Figure 3c), indicating that filament formation is effectively irreversible on the experimental timescale.

### Aggregates formed by 1P_352_ transformed into nanoscopic filaments after 10 days

In cells, Tau aggregation occurs over extended time scales, from months to years^31^. To approximate these conditions *in vitro*, we incubated 1P_352_ at 4 for 10 days at a pH of 6.5 (Figure 4a) and used SEM to analyze the resulting filaments. The samples contained entangled filaments with smooth surfaces, consistent with a full conversion of droplets into string-like assemblies, with an average width of 22 ± 3 nm (Figure 4b,c). These findings support a model in which droplets act as seeds for aggregation when the pH falls below 6.8, shrinking and agglomerating to initiate filament formation that remains thermodynamically stable even after the pH is restored.

**Figure 4.**
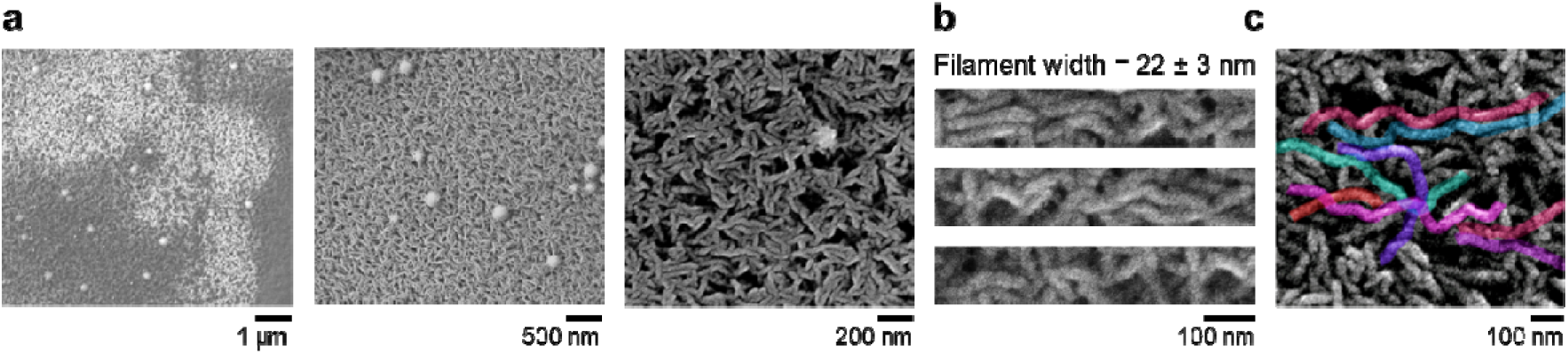
Filaments formed by 1P_352_ after 10 days of incubation. SEM images of the 1P_352_ filaments observed after incubation for 10 days. a) The filaments at increased magnification from left to right. b) Further enlargement of the filaments c) As B, with filaments colored using the Affinity designer software, emphasizing the structure, length and morphology of the filaments.

### Molecular dynamics simulations demonstrate protonation-dependent stabilization of pS352 amyloid structure upon decreasing the pH

The effect of acidification at the molecular level was probed using molecular dynamics (MD) simulations to assess peptide stability in an amyloid context. As the sequence does not contain any histidine residue, S352 is the only residue in this peptide that can be protonated at this pH range. pS352 in 1P_352_ was simulated in its deprotonated state, corresponding to pH 7.1, and in its protonated state, corresponding to pH 6.5. At pH 7.1, conformational changes appeared after ∼50 ns, and the peptide largely lost its ordered structure by 100 ns (Figure 5a). In contrast, at pH 6.5, 1P_352_ retained an amyloid-like conformation throughout the simulation, with no major structural changes at either the monomer or filament level (Figure 5b). The width of the simulated filaments is 2.5 nm, an order of magnitude smaller than the 22 ± 3 nm width measured by SEM (Figure 4b,c).

**Figure 5.**
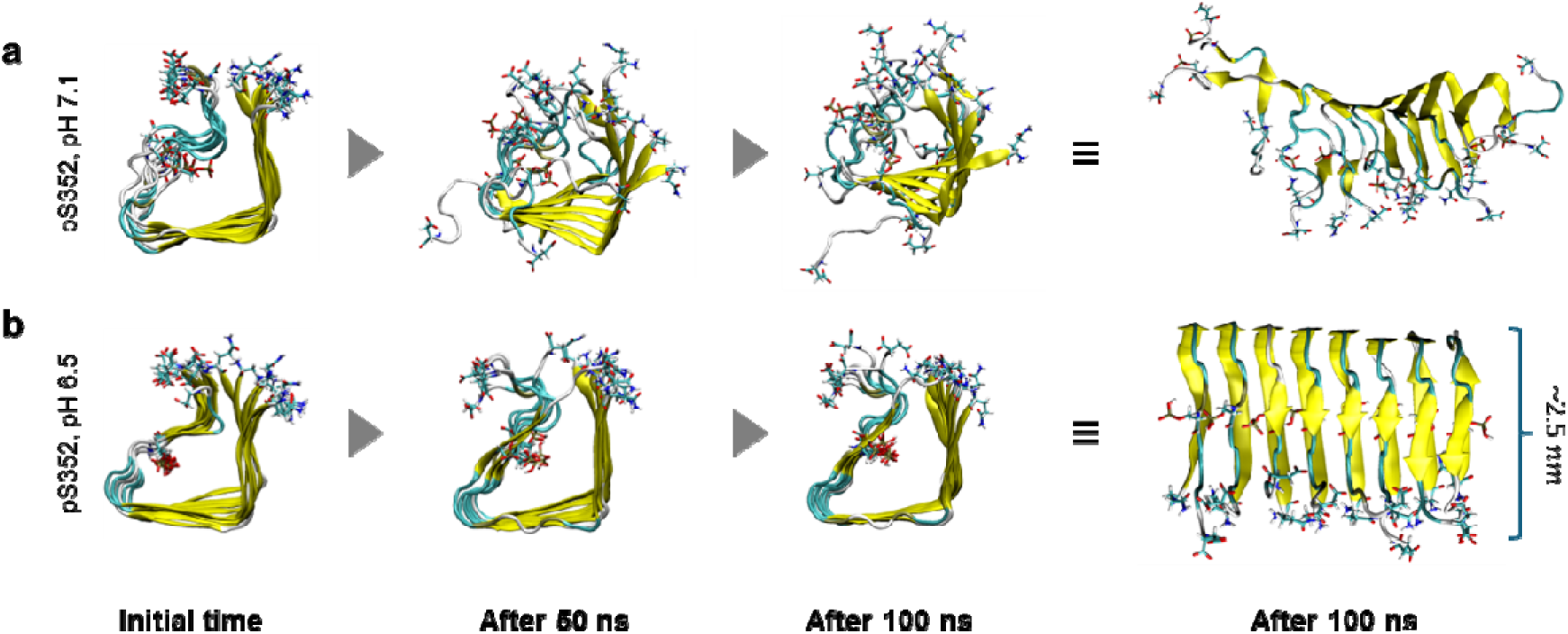
Snapshots from the MD simulations of 1P_352_ reveal pH-dependent stability of amyloid filaments. a) At pH 7.1 (deprotonated pS352), filaments progressively disperse over time; partial disassembly is evident at 50 ns, and most of the filament structure is lost by 100 ns. b) At pH 6.5 (protonated pS352), filaments remain intact, with no major structural changes observed throughout 100 ns of MD simulations.

### pH-dependent protonation of 1P_352_ locally stabilizes the Tau-R4 amyloid core

To investigate the pH-dependent behavior of 1P_352_ at the molecular level, we performed 2D COSY NMR experiments across a pH range of 7.1 to 6.5 in 0.1 pH unit decrements. Peak assignments were made using TOCSY spectra collected at pH 7.1. Among all residues, pSer352 exhibited the most pronounced downfield shift in its amide proton resonance, with a Δδ_HN_ of 28 Hz, indicative of enhanced proton exchange under acidic conditions (Figure 6a). Analysis of the non-exchangeable Hα chemical shift changes (Δδ_Hα_ = δ_Hαfinal_ – δ_Hαinitial_) revealed a clear transition centered at pH 6.81□±□0.02 for pSer352 (Figure 6b).

**Figure 6.**
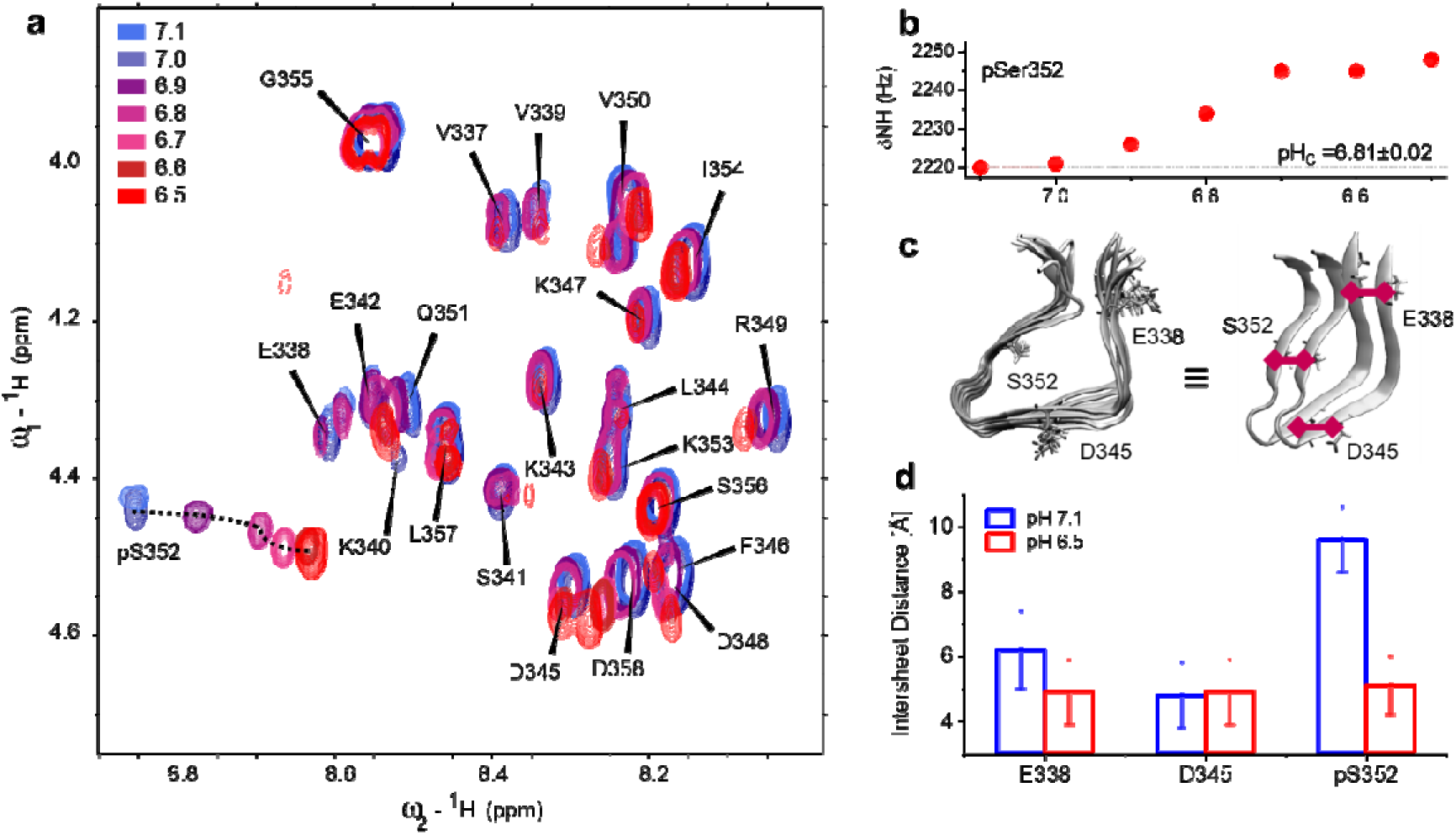
Protonation of pSer352 selectively alters inter-filament spacing in 1P_352_: NMR and MD simulations. a) COSY spectra of 1P_352_ under different pH conditions, ranging from 7.1 (blue) to 6.5 (red). pH values of 7.1-7.0 are shown in blue and transform to purple, dark red, red, pink and light pink as the pH decreases. Each peak is shown with assignment to its residue. The shifts of the peaks of pSer352 are shown by a dotted line. b) The frequency of the NH proton resonance of the phosphoserine as a function of pH. Fitting the δ_NH_ of pSer352 to a sigmoidal function reveals a critical pH of ∼6.8. c) MD-derived structural model of two layers from 1P_352_ amyloid, highlighting the distances between Glu338, Asp345, and Ser352 (red). d) Distances between the above residues in the amyloid state obtained from molecular analysis of MD at pH 7.1 (blue) and pH 6.5 (red).

MD simulations and analyses showed that the peptide adopts a hairpin-like conformation with a hydrophobic core. In this arrangement, the phosphoserine at position 352 points toward the interior of the core (Figure 6c). The inter-sheet distance analysis showed that decreasing the pH from 7.1 to 6.5 did not led to a substantial change in the distance between the two peptide layers at Glu338 or Asp345. In contrast, the distance between the two pSer352 residues decreased markedly, from about 1 nm to about 0.5 nm upon protonation (Figure 6d). These results suggest that filament stability is regulated by local pH effects at pSer352 rather than by structural changes in other regions of the peptide.

### Acidic pH stabilizes a **β**-hairpin centered on pSer352 in Tau-R4 filaments

NMR spectroscopy was used to relate the local conformation of 1P_352_ to its pH-dependent aggregation. NOESY spectra (Figure 7a) show that residues Leu344 to Val350 form a loop, with a dense network of non-HN-Hα*_(i, i-1)_* side-chain NOEs centered on residues 346-350. For example, K347 has Hβ NOEs to F346; D348 shows Hβ/Hγ NOEs to K347; R349 shows Hβ and Hγ NOEs to D348 and V350; D345 shows a tentative Hβ NOE to L344; Q351 exhibits a side-chain NOE consistent with V347 Hγ or L357 Hδ; and K343 shows an Hβ NOE to V350. These contacts cluster around residues 346-350 and support close short-range *i, i±1* packing together with a medium-range K343–V350 contact, consistent with a compact turn or loop that positions pSer352 near the tip of the hairpin structure. Secondary-structure propensity analysis along the Tau-R4 sequence (Figure 7b) reveals a central β-structured segment flanked by additional β-prone regions, placing Ser352 near the tip of a β-hairpin.

**Figure 7.**
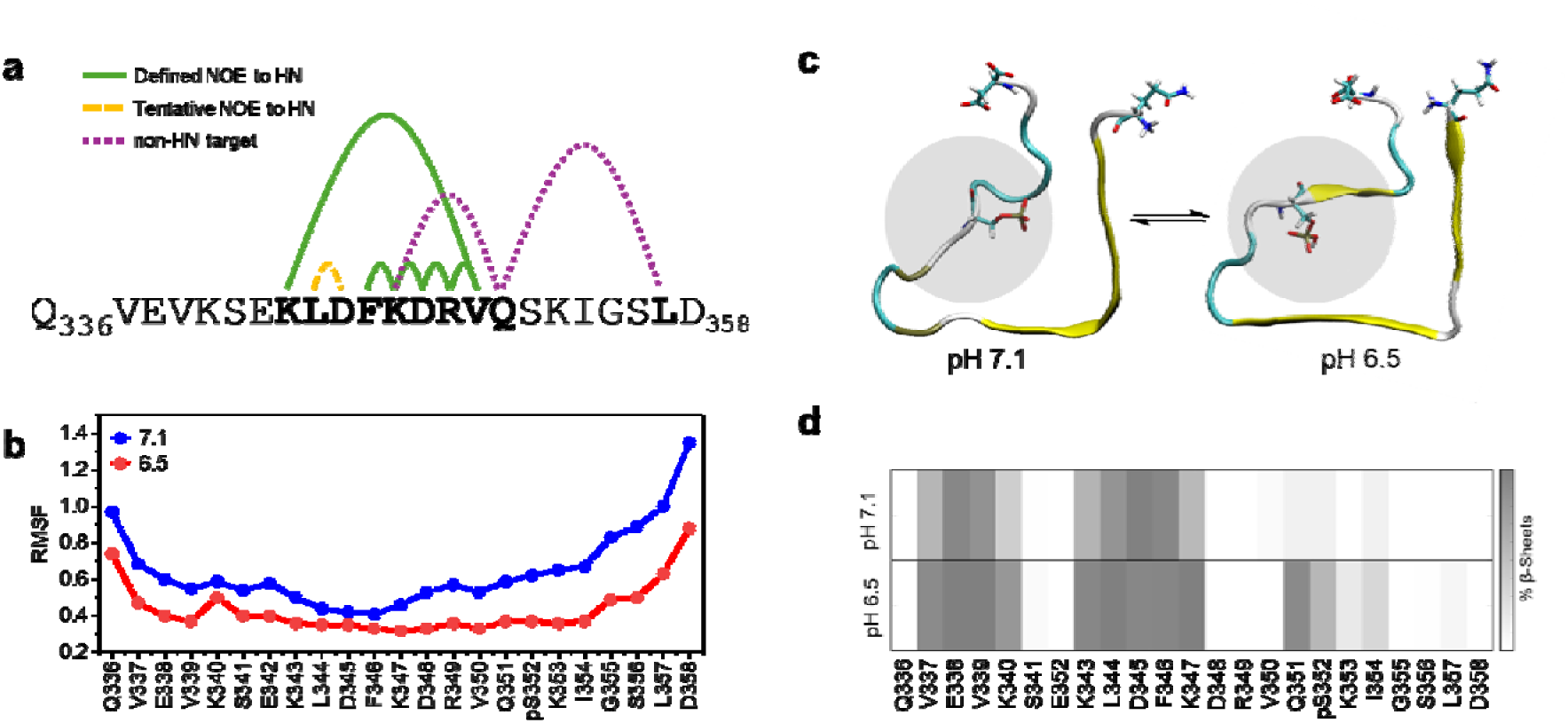
Acidic pH stabilizes local β-structure around pSer352 in Tau-R4 filaments. a) Summary of NOESY side-chain contacts in the central region (Leu344-Val350). Non-HN–Hα*_(i, i±1)_* NOEs cluster around residues 346–350 and include cross-peaks between K347-F346, D348-K347, R349–D348/V350, D345-L344, and K343-V350, defining a K343–V350 loop. A tentative Q351 cross-peak (V347 Hγ or L357 Hδ) suggests additional contacts flanking the loop. b) Root-mean-square fluctuation (RMSF) calculation shows flexibility per residue at the peptide level. The calculation suggests that at pH of 7.1, the C-terminus region of the peptide, starting at residue Lys347, is less structured than at pH of 6.5. c) Structural models of 1P_352_ within the amyloid filament at pH 7.1 and pH 6.5. The peptide adopts a hairpin-like conformation with a compact hydrophobic core, and pSer352 is oriented toward the interior of this core. At pH 6.5 the hairpin is more compact, whereas at pH 7.1 the backbone is more expanded. d) The percentage of β-sheet propensity per residue from MD simulations at pH 7.1 and 6.5. β-structure is globally increased at pH 6.5, with enhanced β-content in the region surrounding Ser352, indicating a more ordered filament at acidic pH.

MD simulations were performed to examine filament models at neutral and acidic pH (Figure 7c-d). In these models, Tau-R4 adopts a hairpin-like conformation with a compact hydrophobic core, in which pSer352 points toward the interior of this core (Figure 7b). At pH 6.5, the hairpin is more compact and well packed, whereas at pH 7.1 the backbone is more expanded and the core less tightly organized. Consistent with this architecture, MD simulations show that the fraction of residues in β-sheet conformation is globally higher at pH 6.5 than at pH 7.1, with a marked increase in β-sheet content around Ser352 under acidic conditions (Figure 7c, d). Together, the NMR data and the MD simulations indicate that acidification stabilizes a more ordered, β-rich filament state of 1P_352_, whereas at neutral pH the same segment is less structured and more susceptible to disruption.

### 31P NMR reveals that pSer352 protonation stabilizes local structure in Tau-R4 filaments

Phosphoserines typically display pK_a_ values in the range of 5.6^28^-6.1^29,30^, which is close to the pH values tested in this study. ^31^P NMR chemical shift in the pH range of 7.1-6.2 was used to determine whether the conformational change at pSer352 is protonation dependent (Figure 8a). The signal moved up-field from 3.15 ppm at pH 7.1 to 1.49 ppm at pH 6.2. The dependence was sigmoidal. The fit exhibited a critical pH at 6.53 ± 0.02, while the transition lies within 0.3 pH units of the Hα midpoint (6.81 ± 0.02) (Figure 8b). This strongly indicated that the observed pK_a_ corresponds to the protonation transition of the phosphoserine group (Figure 8c)^32–35^.

**Figure 8.**
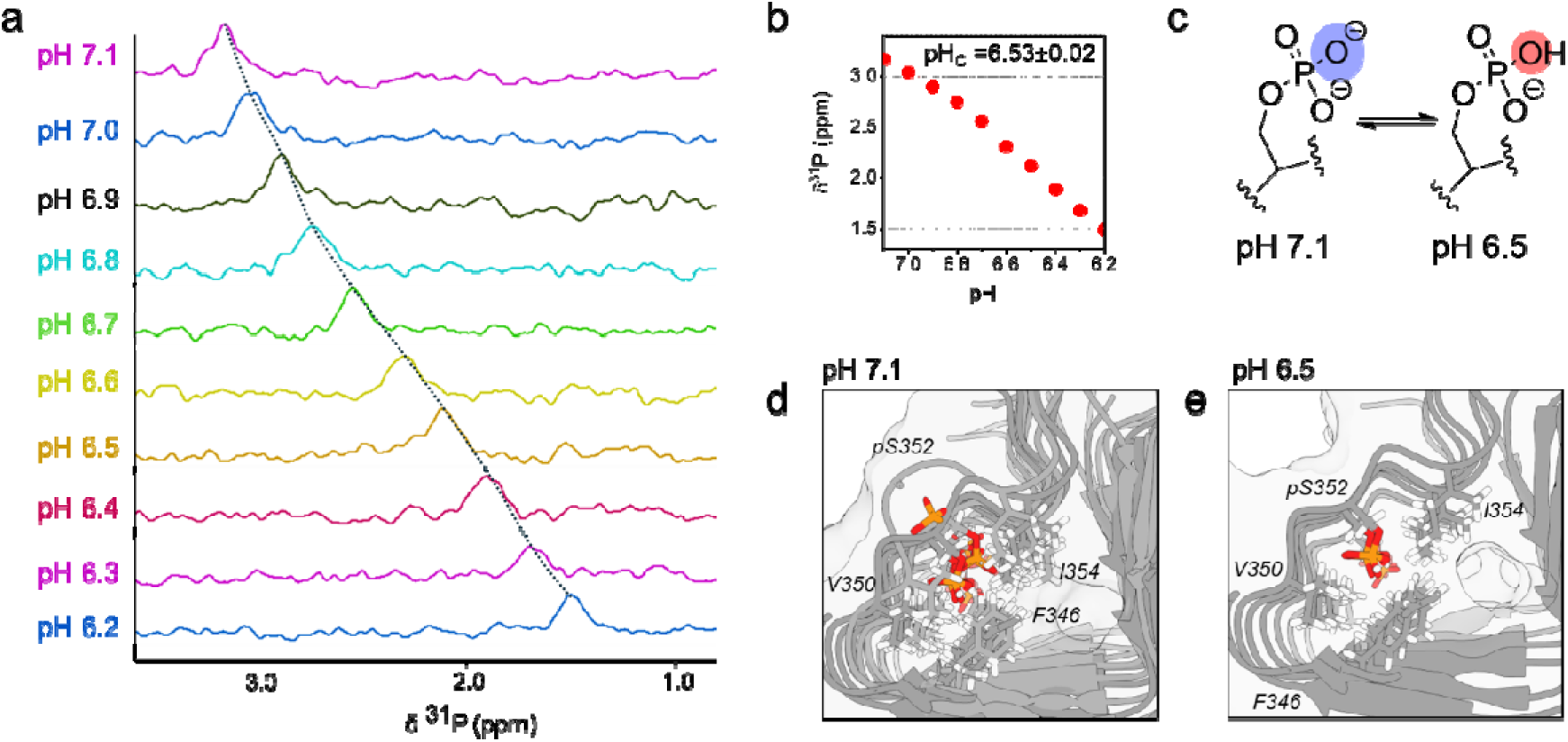
Protonation of pSer352 stabilizes local structure in 1P_352_ filaments. a) ^31^P NMR spectra of 1P_352_ recorded between pH 7.1 and 6.2, showing an upfield shift of the phosphoserine resonance as pH decreases (dashed line). b) Chemical-shift titration of the ^31^P signal (red symbols) fit a sigmoidal model, yielding an apparent pK_a_ of 6.53 ± 0.02. c) Schematic representation of the deprotonated (blue) and protonated (red) states of the phosphate group at pSer352. d) Filament model as simulated by MD, at pH 7.1 highlighting pSer352 and neighboring residues F346, V350 and I354 side chains are more dispersed and the local packing is heterogeneous. e) Corresponding filament structure at pH 6.5, showing tighter and more symmetric packing of F346, V350 and I354 around pSer352, consistent with a more ordered, stabilized local environment in the acidic state.

In the MD-derived structural ensembles (Figure 8d, e), the local packing around pSer352 becomes markedly more ordered at acidic pH. At pH 7.1 (Figure 8d), the side chains of F346, V350 and I354 surrounding pSer352 are more dispersed and the backbone conformations are heterogeneous, indicating a frustrated, less well packed core. At pH 6.5 (Figure 8e), the same residues overlay more tightly and symmetrically around pSer352, consistent with a compact, stabilized local environment within the filament.

## DISCUSSION

In the current study we found that a single site-specific phosphorylation is a trigger that links mild acidification to Tau aggregation in disease. Protonation of pSer352 in a physiologically relevant pH range drives a local structural rearrangement that results in a droplet-to-aggregate transition. This shift yields thin amyloid filaments with increased β-sheet and β-turn structures, providing a direct mechanistic bridge between a specific phosphorylation site and acid-driven Tau aggregation. This may explain the effect of acidification on Tau aggregation in disease.

### Protonation of pSer352 leads to condensation-mediated Tau-R4_(pSer352)_ aggregation

In the current study, we close a mechanistic gap linking site-specific phosphorylation, mild acidification and Tau-R4_(pSer352)_ aggregation via condensation. Most of the phosphorylation patterns studied did not promote aggregation of the corresponding peptides. The peptides 0P, 1P_356_, 2P_341,356_ and 3P underwent condensation in different pH values, while 1P_341_, 2P_341,352_ and 2P_352,356_ did not undergo condensation or aggregation (Figure 1b). Single protonation of pSer352 is unique, as it allows the Tau-R4 domain to adopt a more compact conformation, which in turn promoted filament formation (Figure 9).

**Figure 9.**
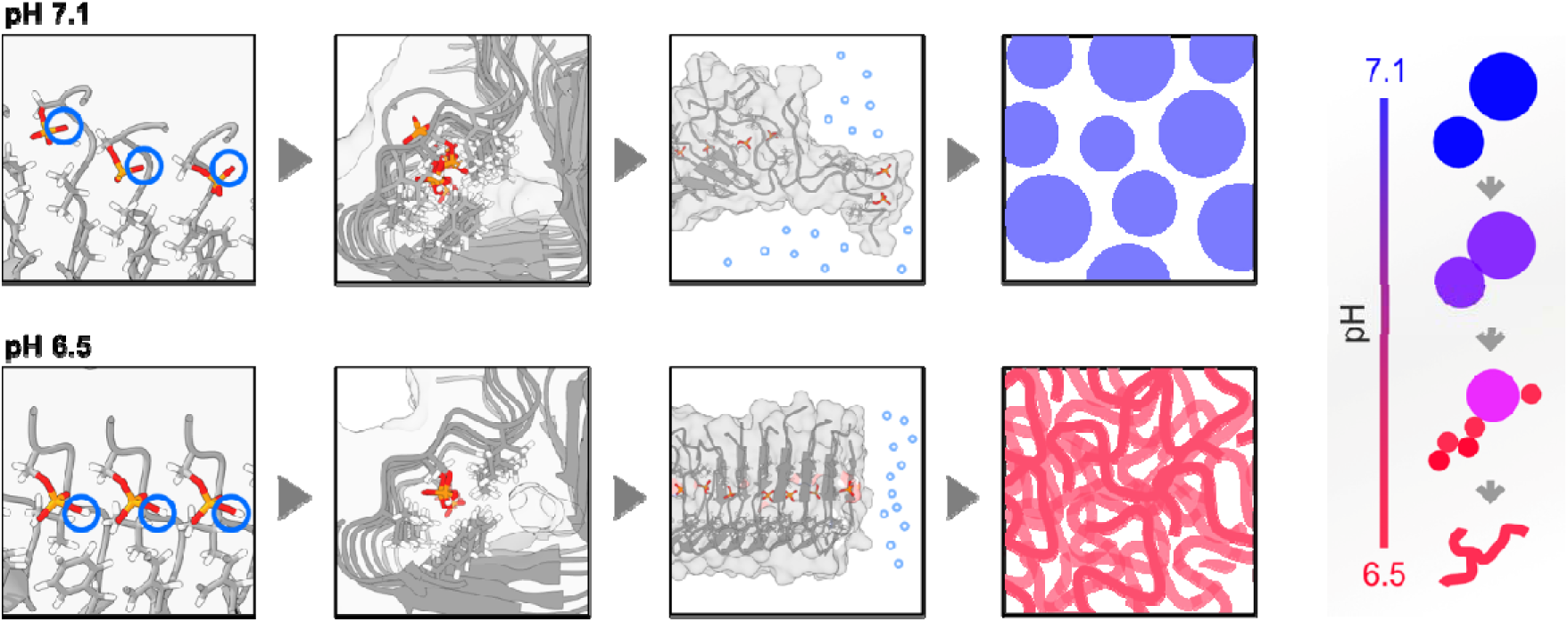
Schematic model for the pH-dependent self-assembly mechanisms of 1P_352_. Top row: At neutral pH 7.1, pSer352 is deprotonated and negatively charged. In this state, the local hairpin around Ser352 is more expanded, packing in the hydrophobic core is loose, and filaments are relatively separated. Under these conditions 1P_352_ forms large, dynamic liquid-like condensates. Bottom row: At mildly acidic pH 6.5, pSer352 becomes partially protonated. Protonation allows tighter local packing around Ser352, leading to a more compact hairpin and smaller inter-filament spacing within the amyloid core. This structural stabilization shifts the peptide from condensates to aggregates. The right panel summarizes the transition along the pH axis: as pH decreases from 7.1 to 6.5, droplets shrink, intermediates with mixed droplets and early filaments appear, and finally a network of entangled filaments is formed.

The first step in this pathway is therefore protonation of the phosphoserine side chain, which we directly monitored by ^31^P NMR. These results are correlated with the idea that specific phosphorylation patterns can support fibril formation, while other do not. Among the library variants, only 1P_352_ undergoes the full condensation-to-aggregate transition upon mild acidification (Figure 1b). The remaining phosphorylation patterns fall into two categories: peptides that condense but do not aggregate (0P, 1P_356_, 2P_341,356_, 3P), and peptides that neither condense nor aggregate (1P_341_, 2P_341,352_, 2P_352,356_)^25^. In the first group, phosphorylation occurs at positions that face the solvent-exposed exterior of the hairpin, where the added charge can promote intermolecular electrostatic interactions sufficient for condensation but does not alter core packing enough to drive a liquid-to-solid transition (Figure 10). In the second group, dual phosphorylation involving pSer352 paired with a second site, likely introduces excessive electrostatic repulsion or steric disruption that destabilizes the hairpin fold altogether, preventing even condensation. The behavior of pSer352 is therefore structurally distinct from all other phosphorylation sites in R4. Because Ser352 sits at the core of the β-hairpin, its side chain is directed into the interior of the hydrophobic core (Figure 7c). This inward orientation means that protonation of pSer352 directly modulates core packing, decreasing the inter-sheet distance at this position from ∼1 nm to ∼0.5 nm (Figure 6d), an effect that no other R4 phosphosite can produce from its solvent-exposed position. This localized compaction is consistent with the increased β-sheet content observed around Ser352 at acidic pH (Figure 7d) and with the preservation of amyloid-like structure throughout MD simulations at pH 6.5 (Figure 5b). Thus, phosphorylation per se is not sufficient to support fibril formation. only phosphorylation at the structurally important position of Ser352, where the hairpin geometry places the phosphate within the core, enables the protonation-driven condensate-to-filament transition (Figure 1b).

**Figure 10.**
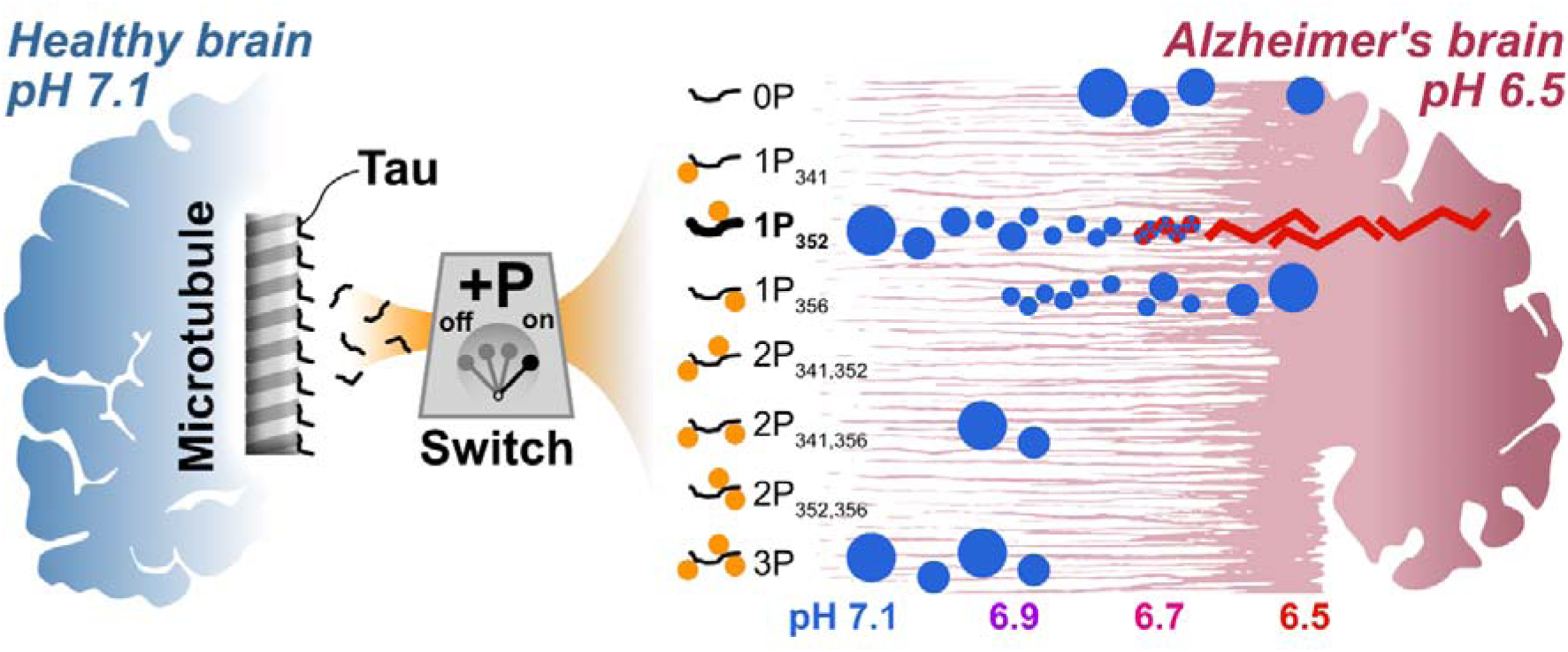
Phosphorylation of Ser352 is a molecular switch for acid-driven Tau-R4 aggregation. In healthy neurons (left, blue), Tau stabilizes microtubules. Hyperphosphorylation detaches Tau, bearing multiple phosphorylation patterns across the R4 domain, from the microtubules. At physiological pH (7.1), R4 with several phosphorylation patterns undergoes condensation (blue circles): 0P forms condensates only at pH 6.7 and below, 1P_356_ and 2P_341,356_ condense at pH 6.9, and 3P condenses at pH 7.1. 1P_341_, 2P_341,352_ and 2P_352,356_ remain soluble at all pH values tested. None of these patterns aggregate upon acidification. Only 1P_352_ undergoes a condensate-to-aggregate transition as the pH decreases from 7.1 to 6.5: condensates shrink, solidify, and convert into amyloid filaments (red). This transition is irreversible and is driven by protonation of pSer352. The resulting filaments accumulate in the Alzheimer’s patients neurons (right, pink). Droplet sizes across all patterns correlate with the phase behavior shown in Figure 1b,c.

^31^P chemical shifts of phosphate groups are well-established reporters of their protonation state: As protonation increases, the ^31^P resonance moves upfield and follows a sigmoidal dependence that obeys the Henderson-Hasselbalch equation (Figure 8a, b). This principle underlies routine intracellular pH measurements from the inorganic phosphate (Pi) peak in ^31^P MRS, where the Pi chemical shift is used as a quantitative pH probe in cells and tissues^36,37^.

For phosphoserine, model peptide titrations give pK_a_ values of ∼ 6.1, with low-pH limits near 0.6 ppm and high-pH limits around 4.9 ppm^29^, and similar titrations of two pSer sites in ovalbumin yield pK_a_ values of 6.00–6.04^30^. In our 1P_352_ titration, the ^31^P resonance of pSer352 shifts over a comparable range, between ∼ 3.5 ppm and ∼ 1.5 ppm, and fits a Henderson-Hasselbalch curve with an apparent pK_a_ of 6.53 ± 0.02. The observed chemical-shift range and pKa are fully consistent with previous phosphoserine titrations, confirming that the ^31^P signal directly reports protonation of pSer352. The lack oh histidine in the sequence means that there are no other residues in sequance with a pK_a_ window of 4-9. This, along with the pK_a_ of phosphate as shown by ^31^P NMR, underscores the role of phosphorylation as a pH-dependent switch for phase seperation.

### Local environment of pSer352 elevates its pK_a_ and allows formation of **β**-hairpin

The elevated effective pK_a_ of pSer352 follows naturally from its local environment. MD places pSer352 toward the interior of a hydrophobic core within a β-hairpin^25^ (Figure 7b-d). NOESY shows a dense network of side-chain NOEs consistent with a compact turn that positions pSer352 near the tip of the hairpin, with a medium-range K343–V350 contact that extends the loop to span residues K343–Q351 (Figure 7a). In such low-dielectric, partially dehydrated environments, charged states are disfavored. This interpretation is in line with previous analyses showing that pSer352 has a lower solvent-accessible surface area than pSer341 and pSer356^25,26^. The deprotonated, doubly charged phosphate is therefore destabilized, shifting the pK_a_ so that protonation occurs at higher pH, within the range of neuronal acidosis. Protonation neutralizes the charge, reduces the desolvation penalty and strengthens local contacts. This effect is site-specific: pSer352 sits in the core of the hairpin, so its protonation directly modifies core packing, whereas other phosphorylation sites in R4 do not occupy an equivalent structural position.

In Alzheimer’s, Tau forms filaments with a core that spans residues V306–F378^26^. Within this core, R4 adopts a compact hairpin-like architecture, constrained by two β-sheets between Lys343 and Ile354. This β-structure is stabilized by hydrophobic clustering and aliphatic/ aromatic side chains, including V339, L344, V350, I354 and F346^26^. Similarly, our results show a dense interaction network at the central domain of R4, between Lys343 and Val350, with short-range NOEs centered on residues 346–350 and a medium-range K343–V350 contact (Figure 7a). Our minimal, cofactor-free 1P_352_ assemblies share key features of this architecture. NOESY contacts define a dense interaction network spanning Lys343–Val350 (Figure 7a). MD simulations, which were run independently of the NMR restraints, show a compact hairpin in which pSer352 is oriented toward a hydrophobic core created by F346, V350 and I354 (Figure 7c, 8e). The convergence of these two independent datasets supports the hairpin geometry. These observations are consistent with the hairpin geometry described for the same R4 region^26^, although our assemblies represent an early-oligomeric state rather than the mature fibril. The 2.5 nm width of the simulated filaments is an order of magnitude smaller than the 22 ± 3 nm width measured by SEM (Figure 4b,c). In the full-length Tau filaments, the disordered N- and C-terminal regions form a fuzzy coat that limits lateral association of protofilaments^26^. In our case, the minimal Tau-R4 peptide lacks these flanking regions. The absence of this steric barrier likely allows individual protofilaments to laterally associate into higher-order bundles, accounting for the larger width observed by SEM. Protonation of pSer352 within this geometry allows tighter local packing across the K343-V350 loop (Figure 7a), promoting β-turn formation and hairpin stabilization.

Our results are also consistent with time-resolved cryo-EM work, showing that Tau filament formation can proceed through polymorphic intermediates. A shared first intermediate amyloid (FIA) was identified during *in vitro* assembly of Tau(297–391), with an ordered core comprising residues 302–316. This FIA is located at the N-terminal part of the MTBR, within the R3 domain, and is therefore outside the sequence window of our R4 construct. Thus, R4 alone does not represent the full folding pathway. Instead, our data support a complementary mechanism: once assembly is initiated elsewhere, for example through an unstable, early intermediate such as the FIA, protonation of pSer352 can provide a site-specific route that stabilizes a compact R4 hairpin and (i) opens an alternative pathway to fibril formation under cofactor-free, mildly acidic conditions, or (ii) provides a thermodynamic stabilization that captures and stabilizes downstream filamentous states that are initiated upstream in the MTBR.

### pSer352 protonation defines a single pH window for local backbone rearrangement and for condensation-mediated aggregation

Backbone and phosphate NMR titrations place the conformational transition of 1P_352_ in a narrow pH window. The Hα chemical-shift titration of pSer352 yields a critical point of 6.81 ± 0.02 (Figure 5b), indicating a well-defined change in its backbone environment. Independently, the ^31^P NMR titration shows the phosphoserine resonance shifting upfield with decreasing pH and fits to an apparent pK_a_ of 6.53 ± 0.02 (Figure 8b). Although ^31^P experiments probe the side chain as opposed to the backbone, the two midpoints lie within 0.3 pH units, consistent with a single protonation event at pSer352 that reorganizes the local backbone and side-chain packing in its immediate vicinity.

At the microscopic scale, condensation-mediated aggregation is tuned by pH in the same narrow range. The droplet-diameter titration of 1P_352_ gives a critical pH of 6.80 ± 0.02 (Figure 3a), marking the transition from large, dynamic droplets to smaller, denser condensates and then to filamentous assemblies, as shown by FRAP (Figure 3b). At pH 7.1, 1P_352_ condensates recovered to 75% of initial intensity with a half-time of 2 ± 1 s (Figure 3b), comparable to the ∼70% recovery previously reported for Tau condensates^16^. At pH 6.5, no fluorescence recovery was observed, indicating a liquid-to-solid transition. This is consistent with the time-dependent loss of FRAP recovery previously reported for Tau condensates^38^, and with the dehydration-driven condensate-to-fibril conversion at high ionic strength^39^. the droplet-diameter midpoint of 6.80 ± 0.02 closely matches the Hα transition of pSer352 at 6.81 ± 0.02 (Figure 5b) and the 31P-derived pKa of 6.53 ± 0.02 (Figure 8b). Thus, the onset of droplet shrinkage and filament formation coincides with the protonation window of pSer352, supporting a mechanism in which a single, site-specific protonation event couples local structural rearrangement to the condensate-to-aggregate transition.

### From Neuronal Acidification to Phosphorylated Tau Fibrils: Closing the Mechanistic Gap

Neuronal acidification has long been implicated in neurodegenerative diseases such as Alzheimer’s disease^7,8^. Tau aggregates under acidic conditions but most aggregation studies of Tau at low-pH have used non-phosphorylated Tau fragments. In many cases, the experimental conditions inculded the use of cofactors such as heparin^40^, lipids^41^ or divalent cations^15^. For example, non-phosphorylated Tau 297-391 formed straight filaments at pH values of 6 and 5 in the presence of MgCl□^15^. In another study, RNA41-mediated aggregation of full-length Tau at pH 6.0 yielded compact cross-β fibrils, demonstrating that Tau forms amyloids at mildly acidic pH in the presence of polyanionic cofactors^42^. In the current study, however, we describe a cofactor-free Tau system in which a naturally-occurring phosphorylation provides the trigger for aggregation. We found that under these conditions, the non-phosphorylated Tau-R4_(336-358)_ domain does not aggregate. Thus, it appears that the experimental conditions can strongly affect the way Tau assembles. Mechanistic studies of condensation of Tau have similarly been conducted mostly with recombinant, non-phosphorylated MTBR-derived fragments such as K18 Tau_(244-372)_^43^. pH was usually explored in condensation studies of Tau as a background variable rather than as a systematically controlled parameter. For instance, the pH dependence of condensation has been probed across several discrete pH values, such as 4.8, 6.8, 7.4, and 8.8, but these experiments did not establish how a mild pH decrease drives condensate maturation into amyloid in a site-defined manner^43^. Studies to determine the mechanism of non-phosphorytlated Tau condensation were performed in a pH range of 7.0-8.8, with the lowest pH examined being 7.0^43–45^. Across these studies, condensation of Tau was found to be driven by electrostatic interactions of the microtuble binding region.

Tau undergoes LLPS across a range of conditions. In low-salt, LLPS is largely driven by intermolecular electrostatic attractions between the acidic N-terminal region and the basic middle to C-terminal domains, with hydrophobic interactions playing a comparatively minor role^46^. Due the dependency of these interactions on charge, modest pH shifts are expected to tune Tau LLPS by altering the protonation state, and thus the charge itself. accordingly, pH is among the physico-chemical parameters reported to modulate Tau phase seperation together with ionic strength and post-translational modifications^47^.

Tau can also unsergo phase separation that is derived by hydrophobic interactions at high ionic strength, where droplets dehydrate, undergo maturation, and can directly yield canonical Tau fibrils^43^. Phosphorylation adds negative charges, and was shown to regulate condensate material properties and liquid-to-solid transitions even though it is not strictly required for LLPS per se^46^. Mild acidification results in a decrease of the effective charge in a site- and pattern-defined manner. Protonation of pS352 neutralizes one phosphate charge within a partially buried R4 hairpin environment. This decreases the energetic penalty of placing a charged phosphate in a low-dielectric environment, and enables tighter packing and beta-turn stabilization, thereby unlocking the condensate-to-aggregate transition. This means that the pH drop does not simply tune condensation globally, but selectively lowers the effective charge at a specific phosphosite to shift the assembly pathway towards the dehydrated, aggregation-prone end of the Tau phase separation/aggregation landscape.

Phosphorylation of Tau can trigger its aggregation^19^, and is tied to weakened neuron communication^48^ and pathalogical implications^49^. Aggregation of Tau can be induced by site-specific phosphorylation in the presence of cofactors. For example, a single phosphorylation at Ser293 or Ser305 of the Tau 441 isoform led to its aggregation^24^ when incubated with heparin, while non-phosphorylated Tau 441 did not aggregate. At pH 7.4, without using cofactors, phosphorylation of Ser341 led to aggregation of Tau-R4, while phosphorylation of Ser352 led to condensation^25^. Here we addressed the critical mechanistic gap: How does a specific phosphorylation event couple to a small, pathologically relevant pH drop to drive a defined phase transition from condensates to fibrils. We show that phosphorylation of Ser352 is the molecular switch for the above acid-driven Tau-R4 aggregation (Figure 10). In healthy neurons, Tau binds microtubules. Hyperphosphorylation releases Tau and generates multiple phosphorylation patterns across the R4 domain, but only 1P352 is competent to convert R4 from condensates to amyloid filaments upon mild acidification, while other patterns either condense without aggregating or remain soluble. Protonation of pSer352 drives this irreversible transition, and the resulting filaments accumulate in Alzheimer’s neurons.. We resolved the above gap using a minimal, cofactor-free Tau-R4 system with a defined phosphorylation and a controlled pH that ranges between the normal and pathological values. In this system, mild acidification promoted condensation of Tau-R4 and Tau-R4_(pS356)_, whereas 1P_352_ undergoes a condensate-to-aggregate transition (Figure 1b, Figure 2). This transition is accompanied by condensate compaction, loss of FRAP recovery, increased β-sheet content, and the emergence of amyloid filaments localized to the Ser352 region (Figure 7).

Our results separate condensation from aggregation within the same physiologically relevant pH window. They also explain why MTBR-containing constructs studied at similar pH values can yield distinct assemblies. In cofactor-rich systems, the MTBR, including R4, can be driven directly toward fibrils. In our cofactor-free R4 system with specific phosphorylation of pS352, the lower pH alone favors condensation and the protonation of a single specific phosphoserine unlocks the pathway to amyloid formation.

In conclusion, the current study reveals that a single protonation of the phosphate group at pS352, which is located in the hydrophobic core of the fibril, allows β-turn formation that enables aggregation at the microscopic level. Biologically, this mechanism provides a direct route by which neuronal acidification could selectively convert phosphorylated Tau condensates into stable pathological amyloid aggregates.

More broadly, our systematic evaluation of all phosphorylation patterns within a controlled, minimal system demonstrates that the functional significance of individual post-translational modifications is not fixed but is instead context-dependent, shaped by the local physicochemical environment. In healthy neurons, where the pH is maintained near 7.1, phosphorylation of Ser352 supports reversible condensation. Under the acidic conditions of the neurodegenerating brain, the same modification becomes a trigger for irreversible amyloid formation. This implies that seeding and aggregation events *in vivo* may shift the hierarchy of PTM contributions, such that phosphorylation sites that are benign or even protective under physiological conditions acquire pathological significance as the cellular environment deteriorates. Systematic studies under controlled conditions, as performed here, are therefore essential for mapping how the interplay between specific PTMs and the local environment determines Tau fate across the continuum from health to disease.

## MATERIALS AND METHODS

### Peptide Synthesis, Labelling, and Purification

Peptides were synthesized on Rink amide resin (0.48 mmol/g loading) after 30 minutes of swelling in DMF. The synthesis employed a stirring-based accelerated peptide synthesis method using a reactor^50^ equipped with a sintered glass filter and a heating jacket connected to a circulating 90 °C water bath. The reactor featured an overhead 5-fin PTFE impeller, with solvents and reagents delivered via a feeding line and drained by vacuum filtration. Each cycle began with Fmoc deprotection, followed by a 5 mL DMF wash, a 1-minute coupling step, and a second wash. Fmoc-Ser(HPO3Bzl)-OH and its subsequent residue were double-coupled. The coupling solution (3 mL) consisted of 3 equivalents of protected amino acid, 2.9 equivalents of HATU, and 8 equivalents of DIEA. Fmoc deprotection was achieved by adding 3 mL of a 0.5% DBU solution in DMF for 10 seconds. Peptides were cleaved using a freshly prepared TFA cocktail and purified using a Waters preparative HPLC with a reverse-phase C18 column and a TDW/ACN gradient. Characterization was performed via electrospray ionization mass spectrometry (ESI-MS). Peptide purity was assessed by analytical HPLC. For FRAP and fluorescence microscopy experiments, peptides were labelled at their N-termini with 5(6)-carboxyfluorescein^25^.

### Fluorescence microscopy

Samples of all peptides (each containing 5 % FL-peptide) at concentration of 2 µM-2 mM in 20 mM condensation buffer (containing phosphate buffer, pH of 7.1, 0.001 % to 10 % PEG, adjusted to ionic strength of 150 mM using NaCl), were analyzed using fluorescence microscopy. Decrease in pH was performed by titration with 40 mM HCl solution, increasing the ionic strength in 5.5 mM, from 150 mM to 155.5 mM. A 10 µL aliquot of the 50 µM peptide solution from each sample was applied to a glass slide. Images were captured using a Zeiss Axio Scope A1 microscope equipped with a green-fluorescent filter and an AxioCam ICc 3 camera. Data acquisition and processing were performed using Zen software. The diameters of the droplets were analyzed using ImageJ 1.54j. the critical pH was calculated with sigmoidal fit in Origin 2024 using the SLogistic3 function 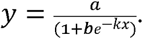 Here, y is the droplet diameter in microns, and x is the pH of the solution. In this SLogistic3 form, a is the fitted upper-plateau diameter (y approaches a as x increases), k (units of 1/pH) is the slope of the transition, and b is a dimensionless position parameter. The critical pH is given by: 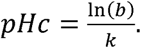

### pH titration and titration cycles for stabilization studies

pS352 was prepared at 50 µM in the buffer described above. A 10 µL sample was applied to a glass slide and imaged as above. The pH was titrated from 7.1 to 6.5 in 0.1-unit steps by adding pre-calculated volumes of 40 mM HCl solution. Samples were mixed, allowed to equilibrate, and imaged at each step. For stabilization cycles, the pH was restored directly to 7.1 in a single adjustment by NaOH, and the samples were re-imaged to assess reversibility. The titration was validated by constructing a calibration curve in the same buffer with known HCl additions, using it to determine the required volume for each target pH, and confirming the volumes in three repetitions. Image analysis was performed in ImageJ 1.54j. Overall, the titration increased the ionic strength by 22 mM, from 150.0 to 172 mM.

### Fluorescence recovery after photobleaching (FRAP)

FRAP samples containing 5% or 95% pS352 were prepared in 25 mM phosphate buffer pH 7.1, 150 mM NaCl, at 50 µM peptide. Immediately before imaging, 1% (v/v) PEG-400 was added and the samples were loaded into an 8-well microslide. FRAP was performed on a Nikon A1R+ confocal microscope (ECLIPSE Ti-E, Nikon) with a 40×/1.3 oil Plan Fluor objective. Imaging used the 488 nm laser; emission was collected at 500–540 nm (GaAsP detection). A defined ROI was bleached at 100 % laser power for 120 ms. Images were acquired every 2 s, with several pre-bleach frames to establish baseline. For achieveing the decrease in pH. HCl was titrated to the sample. Photobleaching correction and recovery analyses were performed in OriginPro 2022. Final recovery curves represent the mean of n=6 condensates.

### Scanning Electron microscopy

pS352 was incubated at 4 °C for 10 days. Samples were deposited on clean silicon wafers, air dried, and imaged on a Quanta 200 scanning electron microscope at 5 kV in low-vacuum mode using a secondary electron detector. Filament width was quantified in ImageJ 1.54j from line profiles on straight segments across multiple micrographs.

### NMR experiments

^1^H-NMR experiments were performed on a Bruker AVII 500 MHz spectrometer operating at the proton frequency of 500.13 MHz, using a 5-mm selective probe equipped with a self-shielded xyz-gradient coil at 23 °C. The transmitter frequency was set on the water signal. Gradient-selected correlation spectroscopy (COSY) experiment using phase-sensitive detection with a quadrature filter^51^; total correlation spectroscopy (TOCSY) using the MLEV-17 spin lock sequence (150 ms) with a homospoil water suppression scheme^52^; and nuclear Overhauser spectroscopy (NOESY) using pulsed field gradients and a phase-sensitive acquisition scheme experiments using a mixing time of 70 ms^53^; were acquired using gradients for water saturation under identical conditions^54^. Spectra were processed and analyzed with TopSpin (Bruker Analytische Messtechnik GmbH) and analyzed and presented with NMRFAM SPARKY software^55^. Resonance assignment followed standard sequential assignment methodology^56^. The δ_H_ shift was fit with a sigmoidal fit in Origin 2024 using the SLogistic3 function 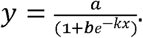 Here y is the observed peak position of the Hα proton (Hz) and x is the solution pH. In this form, a is the fitted upper-plateau value of y, k (units of 1/pH) is the slope, and b is a dimensionless position parameter; the midpoint is given by: *pHc* = 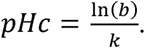

^31^P NMR experiments were performed on a Bruker AVII 500 MHz spectrometer operating at a ^31^P frequency of 202.51 MHz (^1^H frequency 500.27 MHz). One-dimensional ^31^P spectra were acquired using 1024 scans with 2 dummy scans, a spectral width of 49.38 ppm, and an acquisition time of 1.22 s. Chemical shifts were referenced to an external phosphoric acid standard at 0 ppm. The critical pH values were calculated by sigmoidal fitting in Origin 2024 using a logistic function of the form 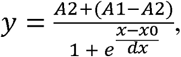 where y is the measured ^31^P chemical shift (Hz) and x is the solution pH. In this form, A1 and A2 are the upper and lower plateaus of y, respectively, x0 is the midpoint (inflection point) corresponding to the half-transition value, and dx (units of pH) is the slope factor that sets the steepness and sign of the transition.

### Constructions of Tau-R4 oligomer models

The phosphorylation on the S352 residue in the Tau-R4 domain was modified in order to fit the determined pH values, thus exploring the effect of protonation states on the phosphorylated R4 domain at the atomic level. Herein, we investigated three fibril-based Tau-R4 domain oligomers, derived from solved cryo-EM (PDB ID code: 5O3T) Tau filaments, the R4 domain ranges between Q336-D358 of the full filament^25^. The three distinct R4 models differ by the phosphorylation state at S352 residue: (1) 7.1 pH model, di-anionic SP2 phosphorylation at S352, with no protonation of the phosphoserine, (2) 6.5 pH model, mono-anionic SP1 phosphorylation with a single protonation. To gain a deeper understanding of the pH effects on the phosphorylated Tau-R4, this project was accompanied by MD simulations of Tau-R4 filament-based oligomers^26^, with models representing different pH conditions of 7.1 and 6.5. A common practice to model MD simulations at different pH values, is to apply the protonation states of any acidic or basic amino acids within the sequence of the protein, which would have passed their pK_a_ or pK_b_ values at the desired pH, thus mimicking the effect of the change of pH on the system^57^. Theoretically, based on the known pK_a_ values found for amino acids in solution, there would have been no such transition of protonation state in the Tau R4 system between these pH values^58^. However, the idea of one single, fixed pK_a_ value has been debated for some time, and the growing consensus is that protein microenvironments significantly influence a residue’s inclination to undergo protonation/de-protonation^59^. It is now recognized that the protonation states of amino acids are a more complex concept than previously conceived, as solution-based pK_a_ values do not account for the effects of the unique protein microenvironment^60,61^. When examining the R4 domain structure, it is apparent that the S352 residue, which undergoes phosphorylation^25^, is in the fibril core and is surrounded by hydrophobic amino acids. This hydrophobic environment destabilizes the charged side chain of the buried phosphorylated serine. Consequently, the protonation from a di-anion to a mono-anion would be preferred at a higher pH than the pK_a_ measured in solution (pK_a_ = 5.70 for phosphoserine in solution)^58^, which could account for protonation between the designated pH values of our system, 7.1 and 6.5. Accordingly, in the MD simulations di-anionic phosphoserine for the pH 7.1 model, and a mono-anionic phosphoserine for the pH 6.5 model were applied and compared to a wild type (WT) model, which was a fibril-like oligomer model of R4 with no phosphorylation. Each phosphorylated Tau R4 oligomer model was constructed by applying the BIOVIA Discovery Studio Visualizer software (BIOVIA, Dassault Systèmes, BIOVIA Discovery Studio, Release 2012, San Diego: Dassault Systèmes, [2023]) and using the CHARMM36 forcefield^62^. Energy minimization was performed using the adopted basis Newton-Raphson (ABNR) method. During the minimization, each S352 was applied with the patch of SP1 and SP2 to form the mono-anionic SP1 and di-anionic SP2 phosphorylations, respectively. The initial structure figure illustrates the minimized phosphorylated Tau-R4 oligomer models. Subsequently, the MD simulations were performed separately for each model.

### Molecular Dynamics (MD) simulations protocol

The MD simulations of the solvated constructed models were performed in the NPT (N-number of particles; P-pressure; T-temperature) ensemble using the nanoscale molecular dynamics (NAMD) package, with the CHARMM36m force-field with the CMAP correlation^62,63^. Each phosphorylated R4 oligomer model was made up of eight identical monomers. The fibril-like oligomers were energy-minimized and explicitly solvated in a TIP3P water box^64,65^. Each water molecule within 2.5 Å of the models was removed. Counter ions (NaCl) were added at random locations to neutralize the charge of the models. The Langevin piston method^62,66,67^ with a decay period of 100 fs and a damping time of 50 fs was used to maintain a constant pressure of 1 atm. A temperature of 330 K was controlled using a Langevin thermostat with a damping coefficient of 10 ps^62^. The short-range van der Waals (VDW) interactions were calculated using the switching function, with a twin range cutoff of 10.0 and 12.0 Å. Long-range electrostatic interactions were calculated using the particle mesh Ewald method with a cutoff of 12.0 Å^68,69^. The equations of motion were integrated using the leapfrog integrator with a step of 1 fs. Counter ions and water molecules were allowed to move. Hydrogen atoms were constrained to equilibrium bonds using the SHAKE algorithm^70^. The minimized solvated systems were energy minimized for 2,000 additional conjugate gradient steps and 20,000 heating steps at 200 K, with all atoms allowed to move. The system was then heated from 200 K to 310 K for 300 ps and equilibrated at 310 K for 300 ps. Simulations were performed for 100 ns for each oligomer model. To approve the timescales of the simulations, root-mean-square-deviation (RMSD) calculations were performed along with the MD simulations (Figure S1). Overall, the simulated models exhibited convergence at the end of approximately 20 ns. Therefore, the timescale of the MD simulations in the current work justifies the choice of timescales. The structures were saved every 10 ps for analysis.

### Structural analyses

The phosphorylated R4 oligomer model structures have been explored by several structural analyses. Conformational changes in the peptide structure were obtained by conducting a RMSD analysis. This allowed us to identify the convergence of each R4 monomer within the dimer along the MD simulations. To evaluate the fluctuations of each residue within the R4 oligomers, root-mean-square fluctuation (RMSF) analysis was performed. To compare the secondary structures of the distinct R4 oligomers, the database of the secondary structure of proteins (DSSP) method was applied^71^. Values of DSSP were calculated along the total time of the simulation for each model. The DSSP method provides the percentage of α-helices or β-strands located along the R4 oligomer sequence. The results of the analyses were averaged over the inner monomers of each model, monomers 2-7, excluding monomers 1 and 8 on the ends of the oligomers. The inter-sheet distances and oligomer diameter were estimated based on the distance between C α of designated residues. For the inter-sheet analyses distances were measured at 3 points between the monomers, one measurement along each of the 3 sides of the peptide: E338, D345 and S352. The diameter was measured for 2 cross-fibril distances, between V337-K347 and K340-S352, for each of the oligomers. The distances were averaged over the inner monomers of each model, as was done for the previous analyses.

## ACKNOWLEDGEMENTS

We would like to thank Dr. Roy Hofmann for his help with the ^31^P NMR experiments. AF thanks the Minerva Center for Bio-Hybrid complex systems and the Saerree K. and Louis P. Fiedler Chair in Chemistry. All of the simulations were performed using the high-performance computational facilities of the Miller lab in the BGU HPC computational center. The support of the BGU HPC computational center staff is greatly appreciated.

## Notes

### Competing Interest Statement

The authors have declared no competing interest.

## REFERENCES

1. Rajmohan, R., and Reddy, P.H. (2017). Amyloid-Beta and Phosphorylated Tau Accumulations Cause Abnormalities at Synapses of Alzheimer’s disease Neurons. J. Alzheimers Dis. 57, 975–999. 10.3233/JAD-160612.

2. Parra Bravo, C., Naguib, S.A., and Gan, L. (2024). Cellular and pathological functions of tau. Nat. Rev. Mol. Cell Biol. 25, 845–864. 10.1038/s41580-024-00753-9.

3. Lee, J.H., Yang, D.S., Goulbourne, C.N., Im, E., Stavrides, P., Pensalfini, A., Chan, H., Bouchet-Marquis, C., Bleiwas, C., Berg, M.J., et al. (2022). Faulty autolysosome acidification in Alzheimer’s disease mouse models induces autophagic build-up of Abeta in neurons, yielding senile plaques. Nat. Neurosci. 25, 688–701. 10.1038/s41593-022-01084-8.

4. Stefani, M., and Dobson, C.M. (2003). Protein aggregation and aggregate toxicity: new insights into protein folding, misfolding diseases and biological evolution. Journal of Molecular Medicine 81, 678–699. 10.1007/s00109-003-0464-5.

5. Basurto-Islas, G., Grundke-Iqbal, I., Tung, Y.C., Liu, F., and Iqbal, K. (2013). Activation of asparaginyl endopeptidase leads to Tau hyperphosphorylation in Alzheimer disease. J. Biol. Chem. 288, 17495–17507. 10.1074/jbc.M112.446070.

6. Zhang, Z., Song, M., Liu, X., Su Kang, S., Duong, D.M., Seyfried, N.T., Cao, X., Cheng, L., Sun, Y.E., Ping Yu, S., et al. (2015). Delta-secretase cleaves amyloid precursor protein and regulates the pathogenesis in Alzheimer’s disease. Nature Communications 6, 8762. 10.1038/ncomms9762.

7. Decker, Y., Németh, E., Schomburg, R., Chemla, A., Fülöp, L., Menger, M.D., Liu, Y., and Fassbender, K. (2021). Decreased pH in the aging brain and Alzheimer’s disease. Neurobiol. Aging 101, 40–49. 10.1016/j.neurobiolaging.2020.12.007.

8. Yates, C.M., Butterworth, J., Tennant, M.C., and Gordon, A. (1990). Enzyme activities in relation to pH and lactate in postmortem brain in Alzheimer-type and other dementias. J. Neurochem. 55, 1624–1630. 10.1111/j.1471-4159.1990.tb04948.x.

9. Lyros, E., Ragoschke-Schumm, A., Kostopoulos, P., Sehr, A., Backens, M., Kalampokini, S., Decker, Y., Lesmeister, M., Liu, Y., Reith, W., and Fassbender, K. (2020). Normal brain aging and Alzheimer’s disease are associated with lower cerebral pH: an in vivo histidine (1)H-MR spectroscopy study. Neurobiol. Aging 87, 60–69. 10.1016/j.neurobiolaging.2019.11.012.

10. Tyrtyshnaia, A.A., Lysenko, L.V., Madamba, F., Manzhulo, I.V., Khotimchenko, M.Y., and Kleschevnikov, A.M. (2016). Acute neuroinflammation provokes intracellular acidification in mouse hippocampus. J. Neuroinflammation 13, 283. 10.1186/s12974-016-0747-8.

11. Amor, S., Puentes, F., Baker, D., and van der Valk, P. (2010). Inflammation in neurodegenerative diseases. Immunology 129, 154–169. 10.1111/j.1365-2567.2009.03225.x.

12. Balestrino, M., and Somjen, G.G. (1988). Concentration of carbon dioxide, interstitial pH and synaptic transmission in hippocampal formation of the rat. J Physiol 396, 247–266. 10.1113/jphysiol.1988.sp016961.

13. Hoyer, W., Antony, T., Cherny, D., Heim, G., Jovin, T.M., and Subramaniam, V. (2002). Dependence of α-Synuclein Aggregate Morphology on Solution Conditions. Journal of Molecular Biology 322, 383–393. 10.1016/S0022-2836(02)00775-1.

14. Wang, H., Wu, J., Sternke-Hoffmann, R., Zheng, W., Mörman, C., and Luo, J. (2022). Multivariate effects of pH, salt, and Zn(2+) ions on Aβ40 fibrillation. Commun. Chem. 5, 171. 10.1038/s42004-022-00786-1.

15. Duan, P., Dregni, A.J., Mammeri, N.E., and Hong, M. (2023). Structure of the nonhelical filament of the Alzheimer’s disease tau core. Proceedings of the National Academy of Sciences 120. 10.1073/pnas.2310067120.

16. Wegmann, S., Eftekharzadeh, B., Tepper, K., Zoltowska, K.M., Bennett, R.E., Dujardin, S., Laskowski, P.R., MacKenzie, D., Kamath, T., Commins, C., et al. (2018). Tau protein liquid-liquid phase separation can initiate tau aggregation. The EMBO Journal 37, e98049. 10.15252/embj.201798049.

17. Eisenberg, D., and Jucker, M. (2012). The Amyloid State of Proteins in Human Diseases. Cell 148, 1188–1203. 10.1016/J.CELL.2012.02.022.

18. Knowles, T.P., Vendruscolo, M., and Dobson, C.M. (2014). The amyloid state and its association with protein misfolding diseases. Nat Rev Mol Cell Biol 15, 384–396. 10.1038/nrm3810.

19. Meng, J.X., Zhang, Y., Saman, D., Haider, A.M., De, S., Sang, J.C., Brown, K., Jiang, K., Humphrey, J., Julian, L., et al. (2022). Hyperphosphorylated tau self-assembles into amorphous aggregates eliciting TLR4-dependent responses. Nature Communications 13, 2692. 10.1038/s41467-022-30461-x.

20. Boyko, S., and Surewicz, W.K. (2022). Tau liquid-liquid phase separation in neurodegenerative diseases. Trends in cell biology 32, 611–623. 10.1016/j.tcb.2022.01.011.

21. Ambadipudi, S., Biernat, J., Riedel, D., Mandelkow, E., and Zweckstetter, M. (2017). Liquid-liquid phase separation of the microtubule-binding repeats of the Alzheimer-related protein Tau. Nature Communications 8, 275. 10.1038/s41467-017-00480-0.

22. Samarasimhareddy, M., Mayer, D., Metanis, N., Veprintsev, D., Hurevich, M., and Friedler, A. (2019). A targeted approach for the synthesis of multi-phosphorylated peptides: a tool for studying the role of phosphorylation patterns in proteins. Organic & Biomolecular Chemistry 17, 9284–9290. 10.1039/c9ob01874c.

23. Bressler, S.G., Mitrany, A., Wenger, A., Nathke, I., and Friedler, A. (2023). The Oligomerization Domains of the APC Protein Mediate Liquid-Liquid Phase Separation That Is Phosphorylation Controlled. Int. J. Mol. Sci. 24, 6478, 6478. 10.3390/ijms24076478.

24. Ellmer, D., Brehs, M., HajlYahya, M., Lashuel, H.A., and Becker, C.F. (2019). Single posttranslational modifications in the central repeat domains of Tau4 impact its aggregation and tubulin binding. Angew. Chem. Int. Ed. 131, 1630–1634. 10.1002/anie.201805238.

25. Bressler, S.G., Grunhaus, D., Aviram, A., Rudiger, S.G.D., Hurevich, M., and Friedler, A. (2025). Specific phosphorylation patterns control the interplay between aggregation and condensation of Tau-R4 peptides. Org Biomol Chem. 10.1039/d5ob00885a.

26. Fitzpatrick, A.W.P., Falcon, B., He, S., Murzin, A.G., Murshudov, G., Garringer, H.J., Crowther, R.A., Ghetti, B., Goedert, M., and Scheres, S.H.W. (2017). Cryo-EM structures of tau filaments from Alzheimer’s disease. Nature 547, 185–190. 10.1038/nature23002.

27. Bressler, Shachar G., Grunhaus, D., Hurevich, M., and Friedler, A. (2026). Methods for studying the effects of phosphorylation patterns in proteins. Biochem. Soc. Trans. 54. 10.1042/bst20250137.

28. Xie, Y., Jiang, Y., and Ben-Amotz, D. (2005). Detection of amino acid and peptide phosphate protonation using Raman spectroscopy. Analytical Biochemistry 343, 223–230. 10.1016/j.ab.2005.05.038.

29. Hoffmann, R., Reichert, I., Wachs, W.O., Zeppezauer, M., and Kalbitzer, H.R. (1994). 1H and 31P NMR spectroscopy of phosphorylated model peptides. Int. J. Pept. Protein Res. 44, 193–198. 10.1111/j.1399-3011.1994.tb00160.x.

30. Vogel, H.J., and Bridger, W.A. (1982). Phosphorus-31 nuclear magnetic resonance studies of the two phosphoserine residues of hen egg white ovalbumin. Biochemistry 21, 5825–5831. 10.1021/bi00266a016.

31. Meisl, G., Hidari, E., Allinson, K., Rittman, T., DeVos, S.L., Sanchez, J.S., Xu, C.K., Duff, K.E., Johnson, K.A., Rowe, J.B., et al. (2021). In vivo rate-determining steps of tau seed accumulation in Alzheimer’s disease. Science Advances 7, eabh1448. 10.1126/sciadv.abh1448.

32. Otvos, J.D., Armitage, I.M., Chlebowski, J.F., and Coleman, J.E. (1979). 31P NMR of alkaline phosphatase. Dependence of phosphate binding stoichiometry on metal ion content. J. Biol. Chem. 254, 4707–4713.

33. Zatina, M.A., Berkowitz, H.D., Gross, G.M., Maris, J.M., and Chance, B. (1986). 31P nuclear magnetic resonance spectroscopy: noninvasive biochemical analysis of the ischemic extremity. J. Vasc. Surg. 3, 411–420. 10.1067/mva.1986.avs0030411.

34. Gillies, R.J., Ugurbil, K., den Hollander, J.A., and Shulman, R.G. (1981). 31P NMR studies of intracellular pH and phosphate metabolism during cell division cycle of Saccharomyces cerevisiae. Proc. Natl. Acad. Sci. USA 78, 2125–2129. 10.1073/pnas.78.4.2125.

35. Grimsley, G.R., Scholtz, J.M., and Pace, C.N. (2009). A summary of the measured pK values of the ionizable groups in folded proteins. Protein Sci 18, 247–251. 10.1002/pro.19.

36. Pantoja-Uceda, D., and Laurents, D. (2025). In Sample pH Measurement by 31P Phosphate NMR. ChemRxiv. 10.26434/chemrxiv-2025-kmpwh.

37. Moon, R.B., and Richards, J.H. (1973). Determination of Intracellular pH by 31P Magnetic Resonance. Journal of Biological Chemistry 248, 7276–7278. 10.1016/s0021-9258(19)43389-9.

38. Hochmair, J., Exner, C., Franck, M., DominguezlBaquero, A., Diez, L., Brognaro, H., Kraushar, M.L., Mielke, T., Radbruch, H., Kaniyappan, S., et al. (2022). Molecular crowding and RNA synergize to promote phase separation, microtubule interaction, and seeding of Tau condensates. The EMBO Journal 41, EMBJ2021108882. 10.15252/embj.2021108882.

39. Tesei, G., Schulze, T.K., Crehuet, R., and Lindorff-Larsen, K. (2021). Accurate model of liquid-liquid phase behavior of intrinsically disordered proteins from optimization of single-chain properties. Proc. Natl. Acad. Sci. USA 118, e2111696118. 10.1073/pnas.2111696118.

40. Zhang, W., Falcon, B., Murzin, A.G., Fan, J., Crowther, R.A., Goedert, M., and Scheres, S.H.W. (2019). Heparin-induced tau filaments are polymorphic and differ from those in Alzheimer’s and Pick’s diseases. eLife 8, e43584. 10.7554/eLife.43584.

41. El Mammeri, N., Gampp, O., Duan, P., and Hong, M. (2023). Membrane-induced tau amyloid fibrils. Commun Biol 6, 467. 10.1038/s42003-023-04847-6.

42. Abskharon, R., Sawaya, M.R., Boyer, D.R., Cao, Q., Nguyen, B.A., Cascio, D., and Eisenberg, D.S. (2022). Cryo-EM structure of RNA-induced tau fibrils reveals a small C-terminal core that may nucleate fibril formation. Proceedings of the National Academy of Sciences 119. 10.1073/pnas.2119952119.

43. Lin, Y., Fichou, Y., Longhini, A.P., Llanes, L.C., Yin, P., Bazan, G.C., Kosik, K.S., and Han, S. (2021). Liquid-Liquid Phase Separation of Tau Driven by Hydrophobic Interaction Facilitates Fibrillization of Tau. J Mol Biol 433, 166731. 10.1016/j.jmb.2020.166731.

44. Najafi, S., Lin, Y., Longhini, A.P., Zhang, X., Delaney, K.T., Kosik, K.S., Fredrickson, G.H., Shea, J.E., and Han, S. (2021). Liquid-liquid phase separation of Tau by self and complex coacervation. Protein Sci 30, 1393–1407. 10.1002/pro.4101.

45. Lin, Y., Fichou, Y., Zeng, Z., Hu, N.Y., and Han, S. (2020). Electrostatically Driven Complex Coacervation and Amyloid Aggregation of Tau Are Independent Processes with Overlapping Conditions. ACS Chem Neurosci 11, 615–627. 10.1021/acschemneuro.9b00627.

46. Boyko, S., Qi, X., Chen, T.-H., Surewicz, K., and Surewicz, W.K. (2019). Liquid–liquid phase separation of tau protein: The crucial role of electrostatic interactions. Journal of Biological Chemistry 294, 11054–11059. 10.1074/jbc.AC119.009198.

47. Rai, S.K., Savastano, A., Singh, P., Mukhopadhyay, S., and Zweckstetter, M. (2021). Liquid-liquid phase separation of tau: From molecular biophysics to physiology and disease. Protein Sci 30, 1294–1314. 10.1002/pro.4093.

48. Watamura, N., Foiani, M.S., Bez, S., Bourdenx, M., Santambrogio, A., Frodsham, C., Camporesi, E., Brinkmalm, G., Zetterberg, H., Patel, S., et al. (2025). In vivo hyperphosphorylation of tau is associated with synaptic loss and behavioral abnormalities in the absence of tau seeds. Nat. Neurosci. 28, 293–307. 10.1038/s41593-024-01829-7.

49. Le, L.T.H.L., Lee, G., Shin, J.W., Shim, Y.-M., Kim, S.-I., Park, S.-H., Won, J.-K., and Lee, M.J. (2025). Phospho-tau Ser356 is mostly confined to pre-NFT neurons in Alzheimer’s pathology. Acta Neuropathologica 150, 62. 10.1007/s00401-025-02967-3.

50. Grunhaus, D., Molina, E.R., Cohen, R., Stein, T., Friedler, A., and Hurevich, M. (2022). Accelerated Multiphosphorylated Peptide Synthesis. Organic Process Research & Development 26, 2492–2497. 10.1021/acs.oprd.2c00164.

51. Aue, W.P., Bartholdi, E., and Ernst, R.R. (1976). Twoldimensional spectroscopy. Application to nuclear magnetic resonance. The Journal of Chemical Physics 64, 2229–2246. 10.1063/1.432450.

52. Bax, A., and Davis, D.G. (1985). MLEV-17-based two-dimensional homonuclear magnetization transfer spectroscopy. Journal of Magnetic Resonance (1969) 65, 355–360. 10.1016/0022-2364(85)90018-6.

53. Kumar, A., Ernst, R.R., and Wüthrich, K. (1980). A two-dimensional nuclear Overhauser enhancement (2D NOE) experiment for the elucidation of complete proton-proton cross-relaxation networks in biological macromolecules. Biochemical and Biophysical Research Communications 95, 1–6. 10.1016/0006-291X(80)90695-6.

54. Liu, M., Mao, X.-a., Ye, C., Huang, H., Nicholson, J.K., and Lindon, J.C. (1998). Improved WATERGATE Pulse Sequences for Solvent Suppression in NMR Spectroscopy. Journal of Magnetic Resonance 132, 125–129. 10.1006/jmre.1998.1405.

55. Lee, W., Tonelli, M., and Markley, J.L. (2014). NMRFAM-SPARKY: enhanced software for biomolecular NMR spectroscopy. Bioinformatics 31, 1325–1327. 10.1093/bioinformatics/btu830.

56. WuDthrich, K. (1986). NMR of proteins and nucleic acids (Wiley).

57. Miller, Y., Ma, B., Tsai, C.-J., and Nussinov, R. (2010). Hollow core of Alzheimer’s Aβ42 amyloid observed by cryoEM is relevant at physiological pH. Proceedings of the National Academy of Sciences 107, 14128–14133. 10.1073/pnas.1004704107.

58. Rumble, J. (2017). CRC handbook of chemistry and physics (CRC press Boca Raton, FL).

59. Porter, M.A., Hall, J.R., Locke, J.C., Jensen, J.H., and Molina, P.A. (2006). Hydrogen bonding is the prime determinant of carboxyl pKa values at the N-termini of α-helices. Proteins: Structure, Function, and Bioinformatics 63, 621–635. 10.1002/prot.20879.

60. Mishra, P., Patni, D., and Jha, S.K. (2021). A pH-dependent protein stability switch coupled to the perturbed pKa of a single ionizable residue. Biophysical Chemistry 274, 106591. 10.1016/j.bpc.2021.106591.

61. Harms, M.J., Castañeda, C.A., Schlessman, J.L., Sue, G.R., Isom, D.G., Cannon, B.R., and García-Moreno E, B. (2009). The pKa Values of Acidic and Basic Residues Buried at the Same Internal Location in a Protein Are Governed by Different Factors. Journal of Molecular Biology 389, 34–47. 10.1016/j.jmb.2009.03.039.

62. Kalé, L., Skeel, R., Bhandarkar, M., Brunner, R., Gursoy, A., Krawetz, N., Phillips, J., Shinozaki, A., Varadarajan, K., and Schulten, K. (1999). NAMD2: Greater Scalability for Parallel Molecular Dynamics. Journal of Computational Physics 151, 283–312. 10.1006/jcph.1999.6201.

63. Huang, J., Rauscher, S., Nawrocki, G., Ran, T., Feig, M., de Groot, B.L., Grubmüller, H., and MacKerell, A.D. (2017). CHARMM36m: an improved force field for folded and intrinsically disordered proteins. Nature Methods 14, 71–73. 10.1038/nmeth.4067.

64. Jorgensen, W.L., Chandrasekhar, J., Madura, J.D., Impey, R.W., and Klein, M.L. (1983). Comparison of simple potential functions for simulating liquid water. The Journal of Chemical Physics 79, 926–935. 10.1063/1.445869.

65. Mahoney, M.W., and Jorgensen, W.L. (2000). A five-site model for liquid water and the reproduction of the density anomaly by rigid, nonpolarizable potential functions. The Journal of Chemical Physics 112, 8910–8922. 10.1063/1.481505.

66. Feller, S.E., Zhang, Y., Pastor, R.W., and Brooks, B.R. (1995). Constant pressure molecular dynamics simulation: The Langevin piston method. The Journal of Chemical Physics 103, 4613–4621. 10.1063/1.470648.

67. Tu, K., Tobias, D.J., and Klein, M.L. (1995). Constant pressure and temperature molecular dynamics simulation of a fully hydrated liquid crystal phase dipalmitoylphosphatidylcholine bilayer. Biophysical Journal 69, 2558–2562. 10.1016/S0006-3495(95)80126-8.

68. Darden, T., York, D., and Pedersen, L. (1993). Particle mesh Ewald: An N⋅ log(N) method for Ewald sums in large systems. The Journal of Chemical Physics 98, 10089–10092. 10.1063/1.464397.

69. Essmann, U., Perera, L., Berkowitz, M.L., Darden, T., Lee, H., and Pedersen, L.G. (1995). A smooth particle mesh Ewald method. The Journal of Chemical Physics 103, 8577–8593. 10.1063/1.470117.

70. Ryckaert, J.-P., Ciccotti, G., and Berendsen, H.J.C. (1977). Numerical integration of the cartesian equations of motion of a system with constraints: molecular dynamics of n-alkanes. Journal of Computational Physics 23, 327–341. 10.1016/0021-9991(77)90098-5.

71. Kabsch, W., and Sander, C. (1983). Dictionary of protein secondary structure: Pattern recognition of hydrogen-bonded and geometrical features. Biopolymers 22, 2577–2637. 10.1002/bip.360221211.

