## Supplementary material for "Site-specific phosphorylation of Ser352 drives aggregation of Tau R4 under acidosis conditions": SI

**Supporting information**

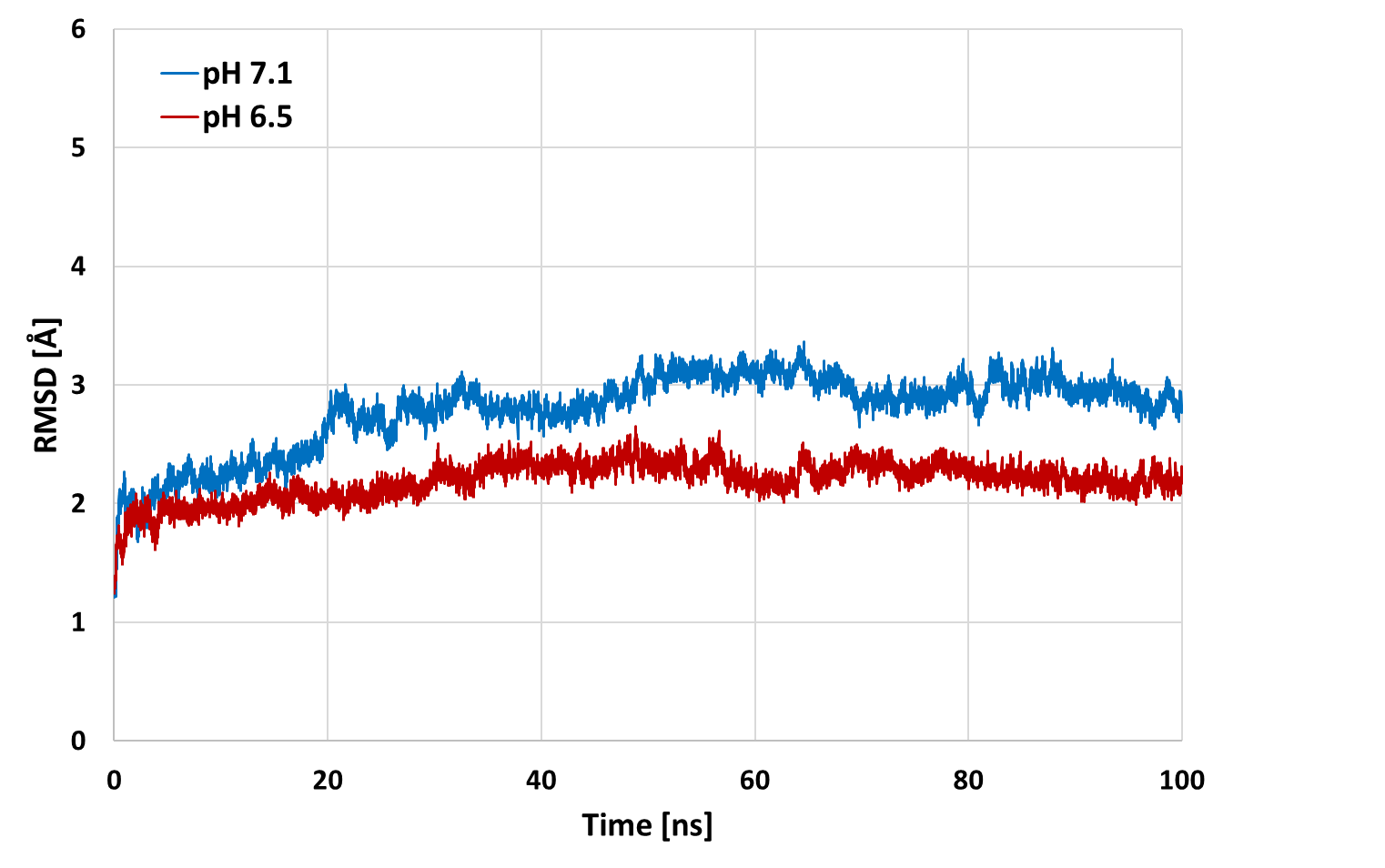

**Figure S1:** Root Mean Square Deviation (RMSD) Values of the pH 7.1 and pH 6.5 1P_352_ Fibril simulations.
